# Mutation of *CFAP57* causes primary ciliary dyskinesia by disrupting the asymmetric targeting of a subset of ciliary inner dynein arms

**DOI:** 10.1101/773028

**Authors:** Ximena M. Bustamante-Marin, Amjad Horani, Mihaela Stoyanova, Wu-Lin Charng, Mathieu Bottier, Patrick R. Sears, Leigh Anne Daniels, Hailey Bowen, Donald F. Conrad, Michael R. Knowles, Lawrence E. Ostrowski, Maimoona A. Zariwala, Susan K. Dutcher

## Abstract

Primary ciliary dyskinesia (PCD) is characterized by chronic airway disease, male infertility, and randomization of the left/right body axis, and is caused by defects of motile cilia and sperm flagella. We screened a cohort of affected individuals that lack an obvious TEM structural phenotype for pathogenic variants using whole exome capture and next generation sequencing. The population sampling probability (PSAP) algorithm identified one subject with a homozygous nonsense variant [(c.1762C>T) p.(Arg588*) exon 11] in the uncharacterized *CFAP57* gene. In normal human nasal epithelial cells, CFAP57 localizes throughout the ciliary axoneme. Analysis of cells from the PCD patient shows a loss of CFAP57, reduced beat frequency, and an alteration in the ciliary waveform. Knockdown of *CFAP57* in human tracheobronchial epithelial cells (hTECs) recapitulates these findings. Phylogenetic analysis showed that CFAP57 is conserved in organisms that assemble motile cilia, and *CFAP57* is allelic with the *BOP2* gene identified previously in *Chlamydomonas*. Two independent, insertional *fap57 Chlamydomonas* mutant strains show reduced swimming velocity and altered waveforms. Tandem mass spectroscopy showed that CFAP57 is missing, and the “g” inner dyneins (DHC7 and DHC3) and the “d” inner dynein (DHC2) are reduced. Our data demonstrate that the FAP57 protein is required for the asymmetric assembly of inner dyneins on only a subset of the microtubule doublets, and this asymmetry is essential for the generation of an effective axonemal waveform. Together, our data identifies mutations in *CFAP57* as a cause of PCD with a specific defect in the inner dynein arm assembly process.

**Significance:** Motile cilia are found throughout eukaryotic organisms and performs essential functions. Primary ciliary dyskinesia (PCD) is a rare disease that affects the function of motile cilia. By applying a novel population sampling probability algorithm (PSAP) that uses large population sequencing databases and pathogenicity prediction algorithms, we identified a variant in an uncharacterized gene, *CFAP57*. This is the first reported example of PCD caused by a mutation that affects only a subset of the inner dynein arms, which are needed to generate the waveform. CFAP57 identifies an address for specific dynein arms. These findings demonstrate the effectiveness of the PSAP algorithm, expand our understanding of the positioning of dynein arms, and identify mutations in *CFAP57* as a cause of PCD.

## Introduction

Motile cilia are complex organelles that project from the surface of cells and are essential for propelling fluids (e.g., in the airways, ventricles) or for providing cell locomotion (e.g., sperm, *Chlamydomonas*). The microtubule-based axoneme shows the classic structure of 9 doublet and 2 central pair microtubules, which is conserved throughout the eukaryotic lineage. Motility comes from the coordinated activity of inner and outer dynein arms (IDA and ODA, respectively) that are attached to the A tubule of the microtubule doublets. Defects in motile cilia cause primary ciliary dyskinesia (PCD), which is a genetically and phenotypically heterogeneous disorder that is characterized by chronic and debilitating respiratory disease, and is frequently accompanied by laterality defects (∼50% of patients) due to abnormal left-right body asymmetry (1). The most common genetic lesions that cause PCD are those that affect components of the ODA, including DNAH5, DNAI1, and DNAH11 (2–4). Another group of PCD associated genes encode ODA-docking complex components (CCDC114 and CCDC151) (5–7), or the proteins that play a role in the cytoplasmic assembly of dynein (SPAG1, DNAAF1-3, HEATR2, and DYX1C1) (8–14). In addition, mutations in two genes, CCNO (15) and MCIDAS (16), cause a PCD-like phenotype by greatly reducing the number of motile cilia. Although genetic variants that cause PCD have been identified in over 40 genes (2, 17–23), there are individuals with confirmed clinical features of PCD but normal axonemal structure as determined by transmission electron microscopy, for whom the genetic basis of their disease is unknown.

In an ongoing effort, we have performed whole exome capture sequencing on more than 400 unrelated cases in order to identify genetic causes of PCD. Within this cohort, there were 99 unrelated cases with clinical features of PCD that include chronic oto-sino-pulmonary symptoms, and low levels of nasal nitric oxide who presented with no obvious axonemal defects by transmission electron microscopy (TEM). Using a new algorithm to analyze exome sequencing data (24), we identified an apparently homozygous stop-gain variant in ciliary and flagella associated protein 57 (*CFAP57*; MIM: 614259; NM_152498.3) in a patient with classical symptoms of PCD that included bronchiectasis, neonatal respiratory distress, otitis media, and sinusitis.

*CFAP57* is allelic with the *BOP2* gene identified previously in *Chlamydomonas*. The *BOP2* locus was first identified as a suppressor of the swimming defect of the *pf10* mutant (25). Double mutant analysis, using IDA and ODA mutants, suggested that the mutation in the *BOP2* locus affects only the IDAs. Electron tomography of the *bop2* mutant showed that a subset of dynein arms were missing on only certain microtubule doublets (26). Specifically, electron tomography revealed reduced intensity on doublets 5, 6, and 8, while doublet 9 showed an intermediate loss of intensity (26).

To date, no pathogenic variants have been identified in proteins that affect only the IDA complexes in PCD patients. The IDAs play a crucial role in determining axonemal waveform and are more heterogeneous than the ODAs. For example, *Chlamydomonas reinhardtii* has six single headed IDA (a, b, c, d, e, and g) and one double headed IDA (I1/f) that span the 96 nm repeat, as determined by cyro-EM tomography using mutant strains affecting these proteins (27, 28). In addition, there are three unique minor inner dynein arms that are only found in the proximal regions and replace the major IDAs; these include *DHC3*, *DHC4*, and *DHC11* (29). Mutations in IDAs are difficult to detect by TEM cross sections since there are 7 dynein arms in 96 nm, and TEM sections are often 60 nm deep. Thus, TEM lacks the resolution necessary to detect changes in individual IDAs, as a single IDA would be obscured by the other IDAs in the section. Our data suggests that homozygosity of a pathogenic variant of *CFAP57* causes PCD, likely by a failure to assemble a subset of inner dynein arms, and should be considered a candidate gene in cases of PCD with apparently normal axonemal structure by TEM and significantly reduced CBF.

## Results

### Whole exome capture and sequencing identifies a *CFAP57* pathogenic variant in a PCD subject

We studied a 38 year old male from family UNC-1095 (Fig. 1a). The subject has *situs solitus* with a classical PCD phenotype that includes neonatal respiratory distress, otitis media, sinusitis, bronchiectasis, and a low rate of nasal nitric oxide production (40 nl/min; cut-off 77 nl/min) (2), but no axonemal defect was detected by TEM (Fig. 1b) (30), DNA from the subject, who had been previously screened for mutations in genes known to be associated with PCD, was sequenced using the IDT capture reagent (31). We analyzed the resulting genotype data using population sampling probability (PSAP), a statistical framework for assessing the significance of variants from n=1 cases of rare genetic disease (24) (Table S1). A new candidate gene, *CFAP57*, was identified and confirmed by direct Sanger sequencing (Fig. 1c and 1d).

**Figure 1:**
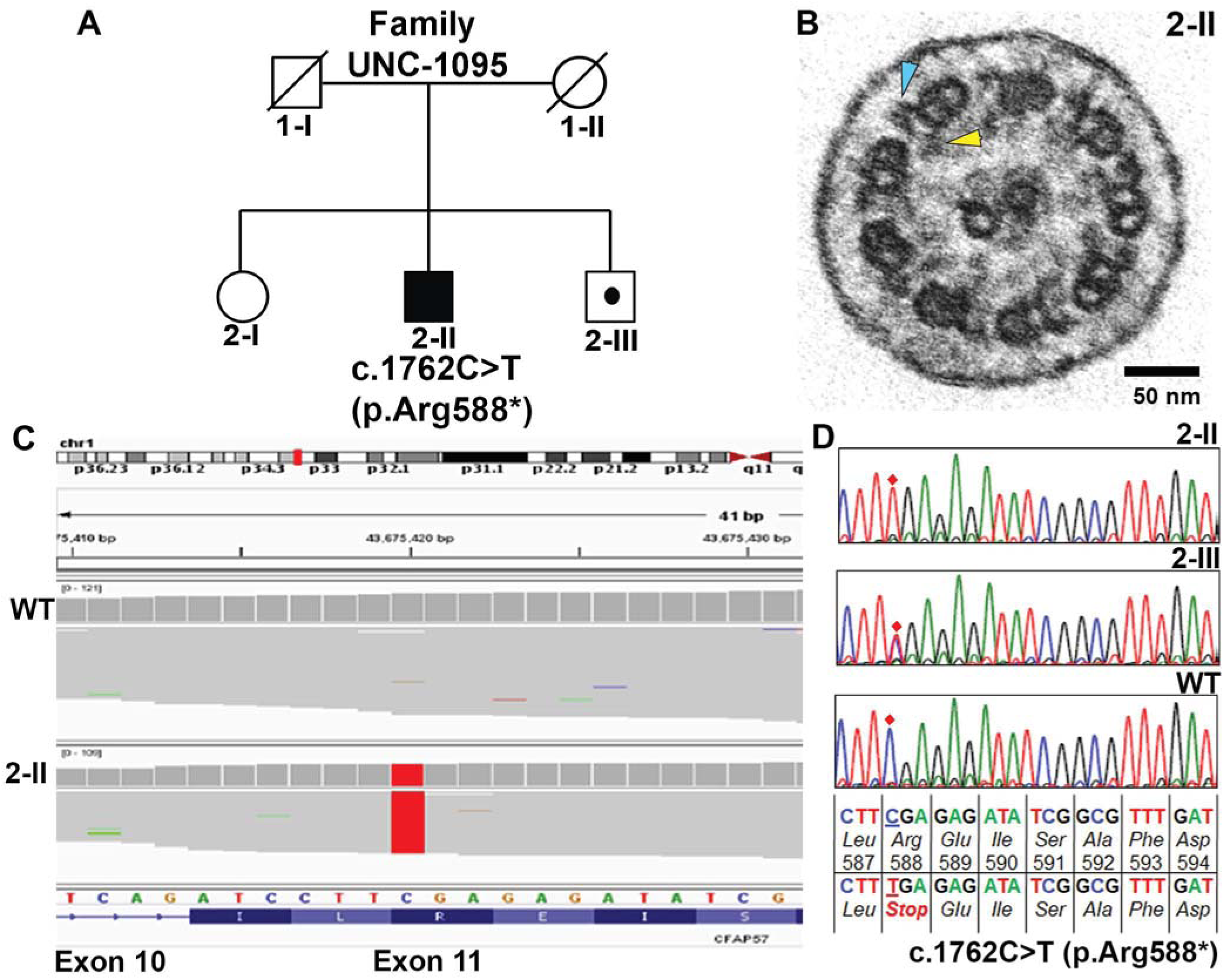
Segregation analysis of pathogenic variant in *CFAP57* found in family UNC-1095. (A) Pedigree analysis of family UNC-1095. Males and females are designated by squares and circles, respectively. Filled symbols indicate PCD affected individual (Proband 2-II). Carrier status is indicated with a dot within the symbols. Symbols with strike-through indicate deceased individual and DNA was not available. (B)Transmission electron micrograph of axonemal cross section of nasal epithelium from proband 2-II showing the central pair surrounded by nine microtubule doublets. Outer dynein arms (blue arrowhead) and inner dynein arms (yellow arrowhead) project from each doublet normally. (C) Whole exome sequencing was carried out for proband 2-II. Snapshot of Bam file showing the location of the apparently homozygous pathogenic variants c.1762C>T (p.Arg588*) in exon 11 of *CFAP57 (WDR65)* NM_001195831.2 on Chr.1p34.2. Position of pathogenic alleles is highlighted in red. (D) Electropherograms showing genomic DNA sequencing. Affected proband (2-II, top panel) is apparently homozygous for c.1762C>T (p.Arg588*) pathogenic allele; whereas, unaffected brother (2-III, middle panel) is a carrier. Base sequence, amino-acid sequence and codon numbers are shown. Position of c.1762C>T base is indicated by red diamond. Base sequence is underlined to indicate the position of pathogenic variants.

Analysis revealed a homozygous stop-gain variant [(c.1762C>T) p.(Arg588*)] in exon 11 of *CFAP57*, which is predicted to result in complete loss of function due to nonsense-mediated decay of the primary *CFAP57* transcripts. Two unaffected siblings were determined to be wild-type and a carrier of the pathogenic variant, consistent with an autosomal pattern of inheritance. CFAP57 is ∼144 kD and contains 10 WD repeats along with 3 predicted coiled-coil domains. PSAP analysis identified the homozygous stop-gain in *CFAP57* as the second most deleterious change in the subject’s genome, following a homozygous missense change in *CD2,* a T-cell surface antigen. Mutations in the latter do not explain the subject’s symptoms (Table S1).

### Expression of *CFAP57* in ciliated human airway epithelial cells

Normal primary human tracheal epithelial cells (hTEC) cells were cultured at an air-liquid interface as previously described (32, 33). Under these conditions, the cells first proliferate as an undifferentiated monolayer and then undergo ciliated cell differentiation. In hTEC cells, the expression of *CFAP57* increased in parallel to the expression of *FOXJ1*, a key gene that drives ciliogenesis (Fig. 2a) and to the expression of *DNAI1*, a known ciliary specific gene (Fig. 2b). Similarly, the levels of CFAP57 protein increased as ciliated airway cells underwent differentiation, as determined by increased levels of FOXJ1 detected by immunoblot (Fig. 2c). This signal was strongly enriched in samples of detergent isolated ciliary axonemes (Fig. 2d). Immunofluorescent staining of CFAP57 in both intact hTEC cultures (Fig. 2e) and isolated cells demonstrate strong positive reactivity throughout the length of the ciliary axoneme (Fig 2f). These results demonstrate that CFAP57 is an axonemal protein, as suggested previously by proteomic analysis of both human and *Chlamydomonas* axonemes (34, 35).

**Figure 2:**
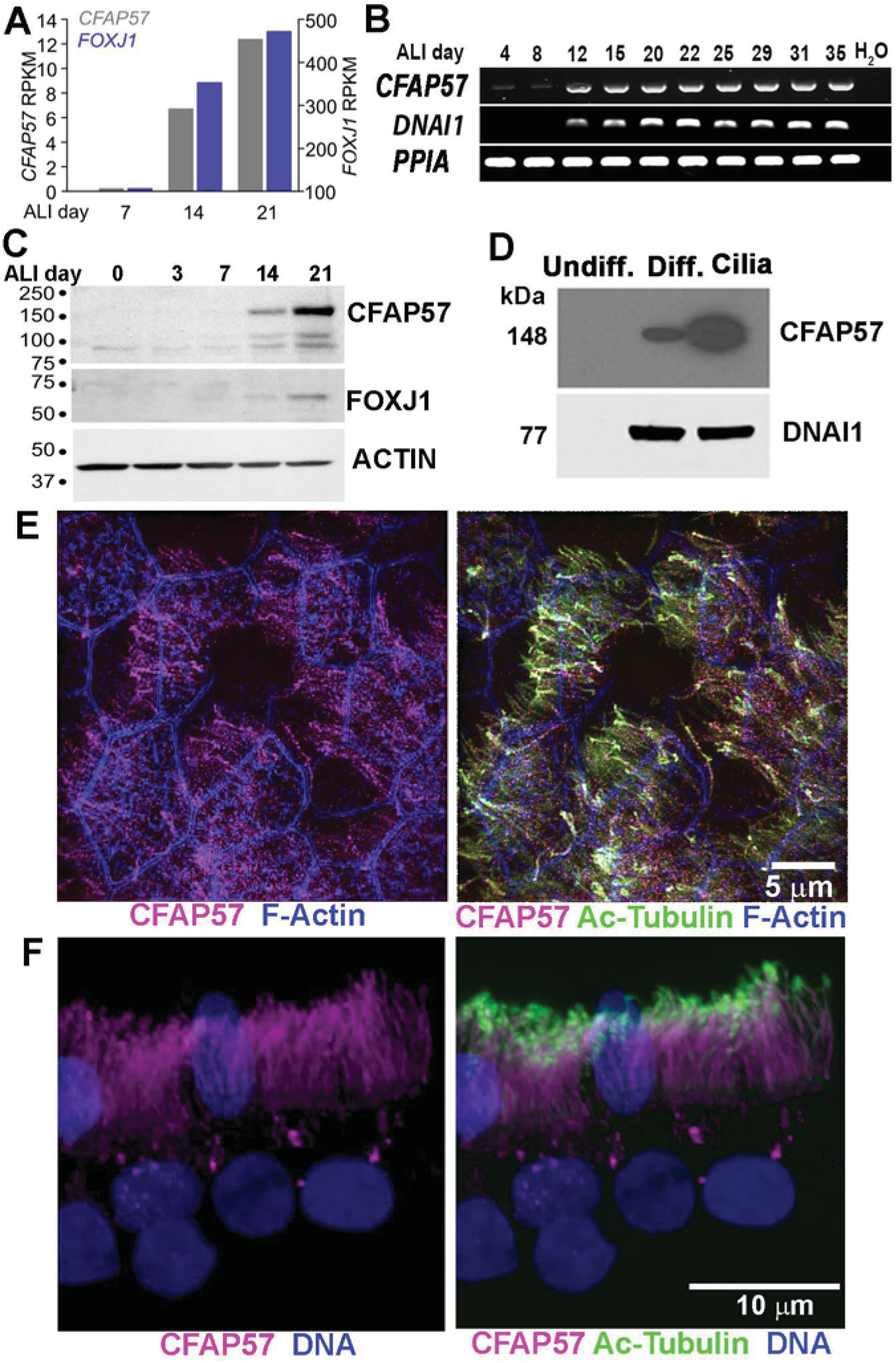
Expression and localization of CFAP57 in human airway epithelial cell. (A and B) Expression of CFAP57 in normal hTEC cells. Total RNA was extracted at the different indicated days of ALI culture. (A) The expression of *CFAP57* correlated with the expression of the expression *FOXJ1*, a gene that drives ciliogenesis and (B) to the expression of *DNAI1,* a ciliated cell specific gene (*PPIA* was used as a housekeeping gene). (C and D) Immunoblots showing (C) increased levels of CFAP57 as ciliated airway cells underwent differentiation (confirmed by increased levels of FOXJ1 levels) and (D) immunoblot showing enrichment of CFAP57 in isolated ciliary axonemes (cilia) compared to differentiated cells (Diff) extract. CFAP57 was absent in protein extract from undifferentiated cells (Undiff.). DNAI1 was use as a loading control for presence of cilia. (E and F) The localization of CFAP57 to the cilia was confirmed by immunofluorescence in (E) whole ALI culture and in (F) isolated ciliated cells.

### A pathogenic variant of *CFAP57* causes defective ciliary beating

To characterize the effects of the genetic variant in *CFAP57* on the function of motile cilia, we obtained human nasal epithelial (HNE) cells from the PCD subject and healthy controls. Cells were expanded in culture as conditionally reprogrammed cells (CRCs) (36), then allowed to differentiate using air-liquid interface cultures as previously described (22). Immunofluorescent staining of isolated ciliated cells from the control cultures showed CFAP57 reactivity along the entire length of the axoneme, while no positive staining was observed in cells from the PCD subject (Fig. 3a). Immunofluorescent intensities for the ODA protein DNAH5 and the radial spoke component RSPH1 were not different between PCD and control cells (Fig. S2), in agreement with the TEM analysis that showed no obvious axonemal structural defect (Fig. 1b). Although the fluorescent intensity of the IDA component, DNALI1, was not obviously different in cells obtained from the PCD subject compared to control cells, it appeared that there was a reduction in intensity at the tip (Fig. 3b; Fig. S1). The lack of CFAP57 resulted in a significant reduction of ciliary beat frequency (CBF) in the PCD cells. Compared to control cells (CBF = 16.98 +/- 4.35 Hz), CBF in the PCD cells was reduced ∼ 30% (10.98 +/- 1.54 Hz) (n= 6; p<0.0007). Using high resolution video microscopy, we performed a detailed analysis of the ciliary waveform. In the PCD cells the waveform appeared slightly altered. In some ciliated cells the waveform appeared normal, while in others, the waveform appeared symmetric, with no clear effective and recovery stroke (Fig. 3c and movies S1, S2). The analysis of maximum displacement from linearity between control (0.53 +/- 0.44 μm) and PCD cells (0.30 +/- 0.10 μm) was not significant (p = 0.3; n=4), indicating that the PCD cells maintained a planar beat. However, the ciliary length of PCD cells (4.7 +/- 0.7μm (s.d.); n=56) compared to control cells (6.0+/- 0.8 (s.d.) μm; n=21) was significantly reduced (p < 0.00001). Thus, the PCD cells exhibited shorter cilia, an altered waveform, and a reduced CBF.

**Figure 3:**
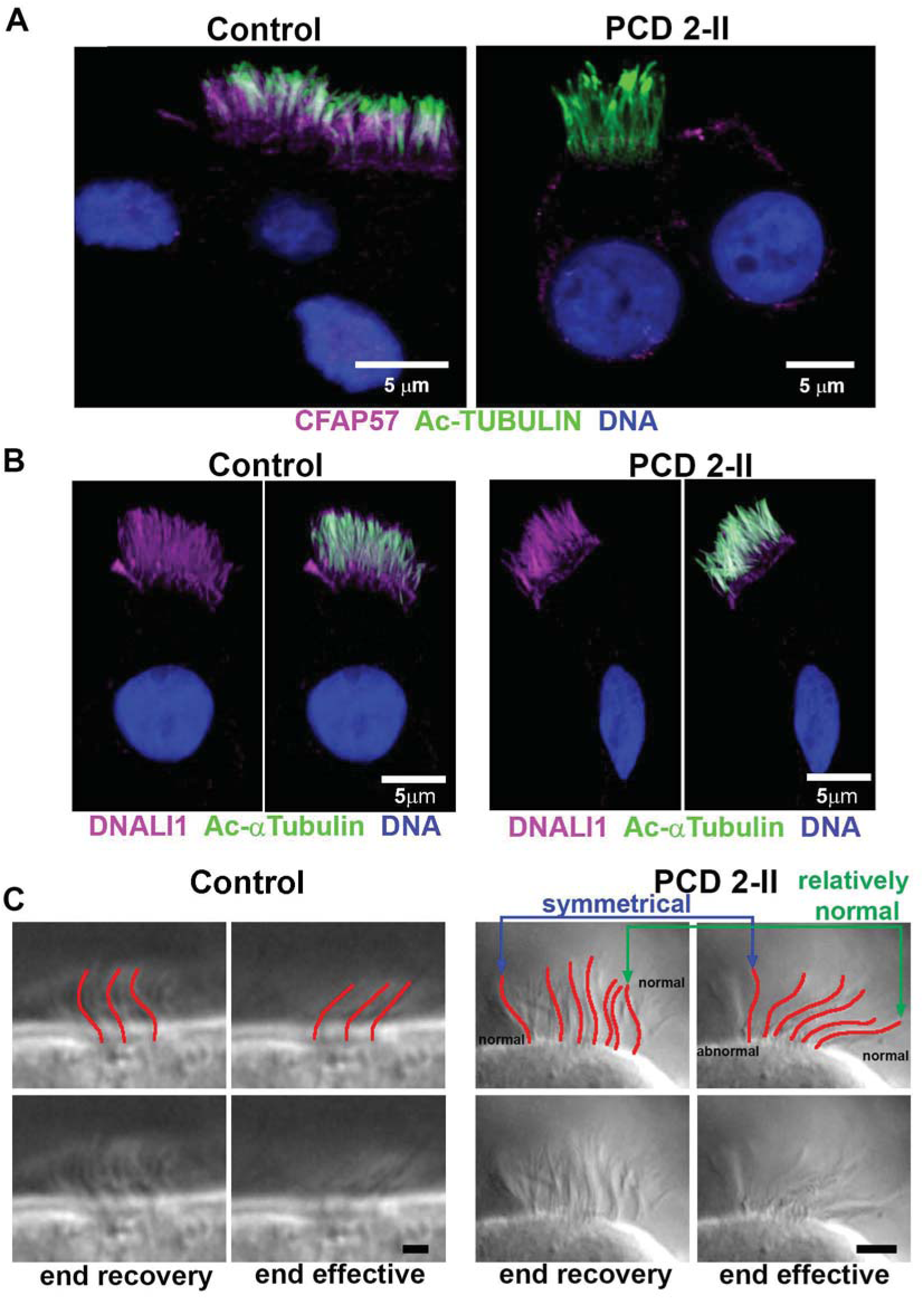
The pathogenic variant in *CFAP57* found in a PCD subject affects ciliary beating. (A) Immunofluorescence of cultured nasal epithelial cells from the PCD 2-II subject (family UNC-1095) and a control (see methods) shows the absence of CFAP57 in ciliated cells from the PCD 2-II subject. (B) DNALI1 immunofluorescent staining of HNE ciliated cells shows a slightly altered localization of the protein. Compared to control cells, in PCD 2-II cells DNALI1 is slightly more abundant at the base of the cilia (apical surface) and is reduced at the tip of the cilia. (C) High-resolution videos of control and PCD 2-II ciliated cells were examined. Compared to control cells, in PCD 2-II cells we observed a heterogeneous waveform. In some cilia the waveform was symmetrical (blue arrows, abnormal) while in other cilia the wave form showed a normal pattern (green arrows). Scale bar = 2 μm.

### *CFAP57* silencing in hTEC results in reduced motile cilia motility

To confirm these results, the role of *CFAP57* in motile cilia function was further defined by silencing expression using an RNAi approach in primary hTEC that were transduced with *a* control plasmid expressing a non-targeted shRNA sequence and a green fluorescent tag (37) or a *CFAP57*-specific shRNA plasmid together with a recombinant lentivirus that contains a cassette that confers puromycin resistance (33). In three biological replicates, *CFAP57* expression was reduced by three out of the four *CFAP57*-specific shRNA sequences when compared to cells transduced with non-targeted shRNA sequences using RT-PCR (Fig 4a). Immunoblot analyses (Fig. 4b) confirmed the absence of the protein from total cell lysates (Fig. 4b), and immunofluorescent staining showed an absence of CFAP57 in ciliated cells. Silencing *CFAP57* did not affect the degree of ciliogenesis in cultured airway cells (Fig.4). High-speed video microscopy analysis of ciliary motility (38) of the *CFAP57*-silenced cultures showed significantly reduced CBF when compared to control cells. (Fig. 4d). Analysis of ciliary beat using high speed video microscopy showed subtle changes in the ciliary waveform, which results in a slightly reduced curvature in *CFAP57*-silenced cells compared to control cells (Movies S3-S6).

**Figure 4.**
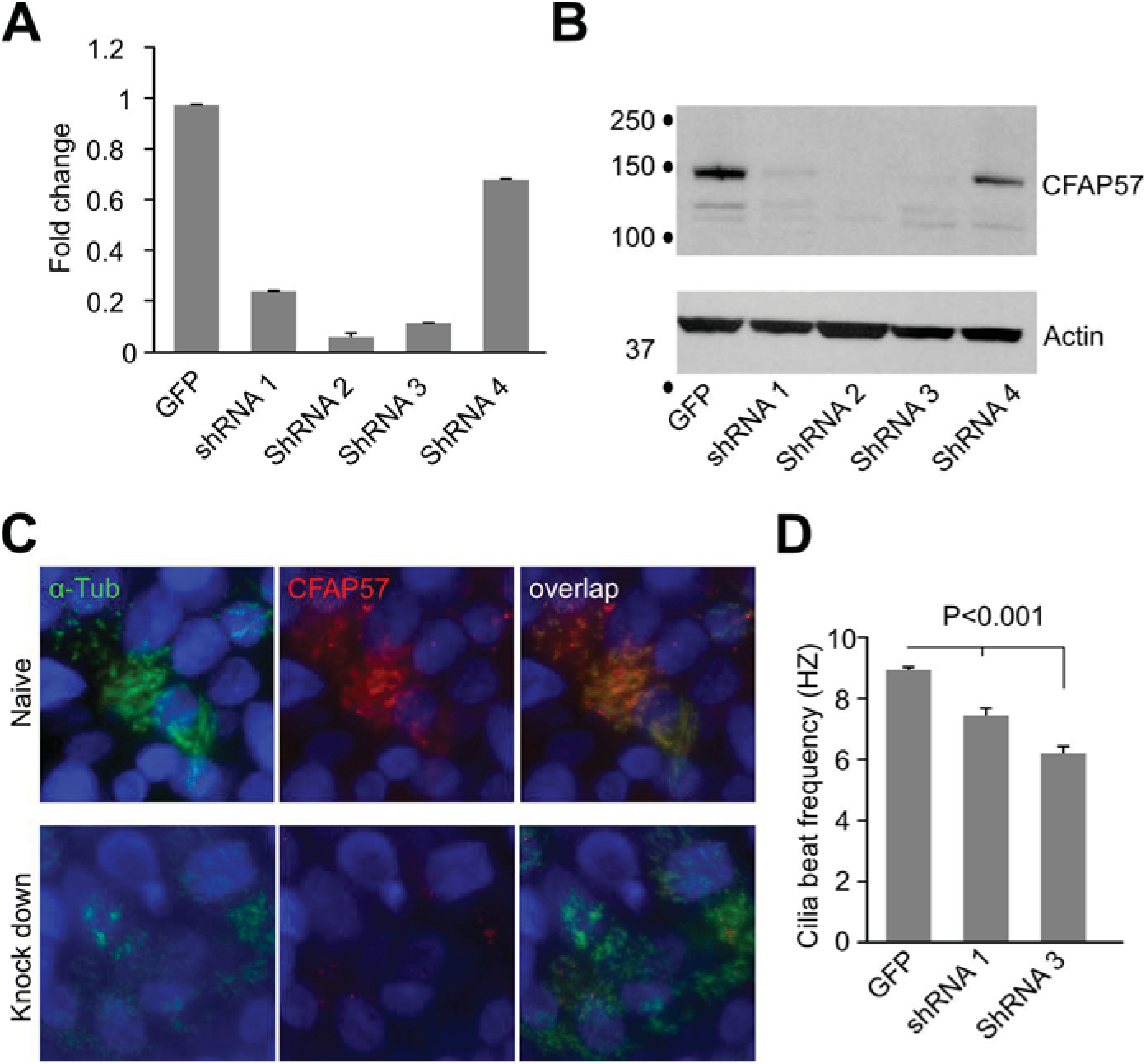
CFAP57 silencing in cultured airway cells causes decreased ciliary motility. (A) qRT-PCR showing fold levels after shRNA specific *CFAP57* silencing in hTEC compared to nontargeted shRNA. (B) Immunoblot analysis of hTEC after shRNA-targeted *CFAP57* silencing. (C) Immunofluorescent staining of hTEC after shRNA-targeted silencing of *CFAP57*. (D) Cilia beat frequency of *CFAP57* silenced cells.

### CFAP57 is conserved in organisms with motile cilia

To evaluate the conservation of CFAP57 across species, we constructed a phylogenetic tree for CFAP57 (Fig. S3). CFAP57 is found in most organisms with motile cilia, except in the cycads, gingkos, and the water fern, *Marselia*. As expected, CFAP57 is missing in organisms that lack motile cilia (flowering plants, nematodes, most fungi). The N-terminus of the protein is predicted to be composed of WD40 repeats that form beta-sheets and the C-terminus is predicted to be -helical (39).

### Effect of *CFAP57* mutations in *Chlamydomonas*

*Chlamydomonas reinhardtii* is an important model for identifying ciliary genes and probing their functions (40). To further investigate the function of CFAP57, we carried out additional studies in *Chlamydomonas*. We obtained three insertional mutant strains (LMJ.RY0402.157050, LMJ.RY402.107706 and LMJ.RY0402.211005) from the CLiP collection, which is an indexed library of insertional mutations in *Chlamydomonas* (41, 42). Insertions in two of the three strains were verified by PCR (Fig. S4). The insertion and drug resistance co-segregate with the swimming phenotype in 21 tetrads for LMJ.RY402.107706 and 7 tetrads for LMJ.RY0402.157050. LMJ.RY402.107706 carries an insert in exon 2 and LMJ.RY0402.157050 carries an insert in intron 7. We examined the mRNA for the two mutants. The transcript in LMJ.RY402.107706 could not be amplified across exons 2 and 3. In strain LMJ.RY402.107050, the transcript for exons 1-7 is present, but we were unable to amplify the remaining transcript (Fig. S4). These alleles failed to complement the *bop2-1* allele in diploid strains and were tightly linked to the locus (n=89 tetrads).

### Defective swimming and ciliary waveform in *fap57 Chlamydomonas*

We analyzed the swimming behavior of these mutants. Swimming velocity analysis shows that *fap57-050* and *fap57-706* swim significantly slower than wild-type cells (CC-125). *fap57-050* was also slower than *fap57-706* (Fig. 5a; Table 1). Ciliary beat frequency extracted from the trajectory was unchanged compared to wild-type cells (Fig. 5b).

**Figure 5.**
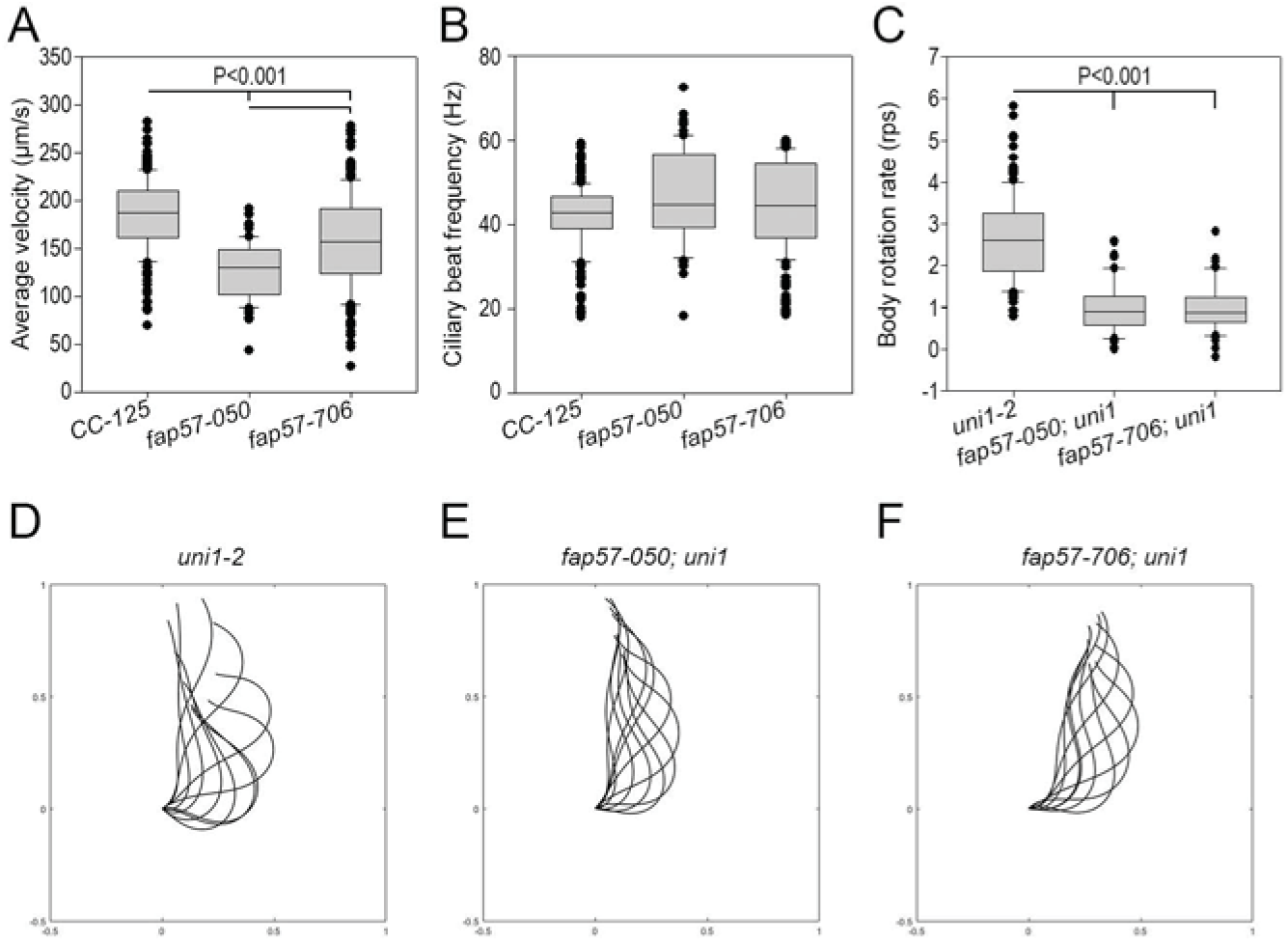
Defective swimming and ciliary waveform in *fap57 Chlamydomonas*. Analysis of swimming behavior of wild-type (CC-125) and *fap57* mutants: (A) Average swimming velocity (μm/s). (B) Ciliary beat frequency (Hz) extracted from the trajectory. Ciliary waveform analysis of uniciliate wild-type (*uni1-2*) and *fap57* mutants. (C) Cell body average rotation rate (revolutions per second, rps). (D) Representative dimensionless (scaled) ciliary waveforms.

**Table 1.** Swimming and ciliary beat frequency of *fap57* mutants

| **Strain** | **Average**<br>**Velocity**<br>**($\mu\text{m/s}$)** | **Ciliary Beat**<br>**Frequency**<br>**(Hz)** | **Cilia length**<br>**($\mu\text{m}$)** | **No. of Cells** |
| --- | --- | --- | --- | --- |
| CC-125<br>(wild-type) | $185.7 \pm 37.8$ | $44.3 \pm 10.3$ | $11.0 \pm 1.9$ | 245 |
| *fap57-050* | $127.9 \pm 29.4$ | $46.9 \pm 10.8$ | $11.1 \pm 2.1$ | 88 |
| *fap57-706* | $157.8 \pm 48.9$ | $44.7 \pm 11.0$ | $10.7 \pm 2.0$ | 205 |

To characterize the waveform of mutants, both the *fap57-050* and *fap57-706* mutants were crossed with the uniciliate mutant *uni1-2* (3). Fifty movies of *fap57-050; uni1* and 51 movies of *fap57-706; uni1* were compared to 153 movies of *uni1-2* (wild-type samples previously described by Bottier *et al.* (44) (Fig. 5a). Body motion analysis shows that both mutants have a significantly slower body rotation rate compared to wild-type cells (Fig. 5c), which agrees with the reduced velocity. Both *fap57* strains have a significantly reduced bend amplitude (Fig. S5b) and an increased average curvature compared to wild-type cells (Fig. S5c). The *fap57-050* mutants also have an average curvature significantly smaller than *fap57-706*, which suggests that the *fap57* mutant waveforms are less wavy, and straighter. Both the amplitude of power and torque are significantly reduced for the *fap57* mutants compared to wild-type (Fig. S4d). Those results are compatible with a reduced swimming velocity as well as a reduced body net rotation. Both alleles displayed waveforms with a shorter stroke.

### Protein composition of *fap57* mutant cilia show a partial phenotype

Using isobaric tags (TMT) for tandem mass spec, we analyzed isolated cilia from four wild-type strains and two technical replicates of two different *fap57* meiotic progeny. There was an average of 107 FAP57 peptides in wild-type cilia, but only 6 peptides in mutant cilia, which suggests a strong loss of function (Table 2). Several of the inner dynein arms (DHC7, 3, 2) are reduced (Table 2) compared to the other dynein heavy chains (Table S3; Fig. S6). The translational elongation factor 1 alpha (EEF1A1) is also reduced in the *fap57* mutant strains to the same extent as the inner dynein arms. This elongation factor is present in proteomic analyses of both *Chlamydomonas* and human axonemes (34, 35, 45–47), which suggests it is not a cytoplasmic contaminant. It can be extracted with KCl and is found in the membrane matrix fraction (34). However, its role in the axoneme is unknown. Four additional proteins are significantly reduced but have few spectral counts (Table 2). Cre12.g540050 and Cre10.g438500 have no orthologs outside of the green algae. Cre13.g562800 has a paralog in *Chlamydomonas* (Cre07.g313850) and both are likely orthologs of WDR49. Cre10.g438500/FAP264 is an ortholog of LRRC74. In humans, WDR49 is expressed in the fallopian tubes and lung and FAM74A is highly expressed in the testes based on the GTEx project (48). They have no known functions. To further examine these additional proteins, we tried a commercial antiserum to WDR49 (Table S5), and it failed to show specific staining in hTECs or *Chlamydomonas*. We examined an insertional mutant strain from the CLiP collection (41) for each of these genes (LMJ.RY0402.139656, LMJ.RY0402.186733, LMJ.RY0402.041374, LMJ.RY0402.255999) but failed to find phenotypes that cosegregated with the insertions.

**Table 2:** TMT proteomics on wild-type and fap57 mutants

| Protein/Gene | No. Peptides | *Fold change relative to all samples* |  |  |
| --- | --- | --- | --- | --- |
|  |  | Average of wild-type controls<br>n=4<sup>a</sup> | *fap57-050*<sup>b</sup> | *fap57-706*<sup>b</sup> |
| FAP57<br>Cre04.g217917 | 107 | 1.0 | 0.1/0.1 | 0.1/0 |
| DHC7; **g**<br>Cre06.g265950 | 341 | 0.98 | 0.5/0.5 | 0.6/0.5 |
| DHC3; minor **g**<br>Cre06.g265950 | 78 | 0.95 | 0.5/0.5 | 0.5/0.6 |
| DHC2; **d**<br>Cre09.g392282 | 540 | 1.0 | 0.8/0.8 | 0.8/0.8 |
| Cre06.g263450<br>Translational<br>elongation<br>factor 1 alpha<br>(EEF1A3) | 220 | 0.98 | 0.3/0.6 | 0.6/0.6 |
| Cre12.g540050 | 5 | 1.05 | 0.3/0.3 | 0.3/0.3 |
| Cre13.g562800<br>Fbox/WD repeat<br>WDR49 | 5 | 1.0 | 0.3/0.3 | 0.3/0.3 |
| Cre03.g174800 | 4 | 0.92 | 0.6/0.7 | 0.5/0.6 |
| Cre10.g438500<br>FAP264<br>FAM74A | 3 | 0.9 | 0.3/0.3 | 0.3/0.2 |

<sup>a</sup> The average of the four axonemal preparations from CC-125 and CC-4533 are averaged.
<sup>b</sup> The ratio for each of the mutant strains is presented

### *CFAP57* mutants suppress the *pf10* mutant

The *CFAP57* gene maps near the *BOP2* locus and the phenotype of the *fap57* mutations is similar to the *bop2* mutant strain. The *bop2-1* mutation results in slow swimming and the loss of inner dynein arms on a subset of the doublets (25, 26). Since *bop2* was isolated as a suppressor of the motility defect of the *pf10* mutant, we tested if the *fap57-706* and *fap57-050* alleles suppress the motility defect of *pf10.* Both alleles acted as suppressors as measured by the number of cells that can oppose gravity in liquid medium (Table S4).

Taken together, the above data demonstrates that CFAP57 plays an important role in the assembly of a specific subset of IDAs, and the proper positioning of these IDAs is important for the generation of normal waveforms. This is consistent with the phenotype observed in the PCD subject, in which absence of *CFAP57* results in reduced CBF and altered waveform, in the absence of a structural defect detectable by TEM.

## Discussion

Mutations in over 40 genes have been reported to cause PCD, and these account for the majority of disease cases (2, 49). However, many cases of PCD (∼30%) remain unsolved at the genetic level. The large number of causative genes are a direct reflection of the complexity of ciliary assembly and structure. Many of the known mutations occur in genes that encode structural proteins of the ODAs or preassembly factors for ODAs and IDAs. To date, no mutations have been reported that cause a defect that affect only the IDAs. We performed exome capture and sequencing on a patient with a clinical diagnosis of PCD who had no known genetic mutation, and no obvious structural changes visible by TEM. Using PSAP, a newly developed algorithm to analyze exome sequence together with large genome databases, we identified a homozygous stop-gain mutation in *CFAP57*. The PSAP algorithm uses large population sequencing databases and pathogenicity prediction algorithms to calculate the probability of sampling a particular genotype, or set of genotypes, observed in a single n=1 case. PSAP is most useful in rare disease studies where families are unavailable, and/or the cohort size is modest (dozens to hundreds). We currently use PSAP on cases of idiopathic nonobstructive azoospermia to identify new causative genes (50), and expect that PSAP will become a useful tool to identify causative genetic variants in other rare diseases. These may include diseases similar to azoospermia and PCD that are characterized by large effect mutations and extensive locus heterogeneity.

In cultured nasal epithelial cells from the PCD patient, we observed a complete absence of CFAP57 by immunofluorescence and the cells exhibited a significantly reduced CBF. Knockdown of *CFAP57* in hTEC cells using shRNA, confirmed by immunofluorescence and immunoblotting, also show a reduced CBF. In contrast, TEM analysis of nasal epithelial cells from the subject appear normal and immunostaining for the ODA protein DNAH5 was unchanged. However, immunostaining for DNALI1 suggested a defect in the localization along the axoneme, with an accumulation at the base and a reduction at the tip. The ortholog of DNAL11 in *Chlamydomonas* is p28/DII1 and it is found in three of the inner dynein arms, including the d dynein. Although the mechanisms that regulate basal CBF have been studied extensively, they remain unclear. These results suggest that a proper balance between ODAs and IDAs is also required to maintain normal CBF. Waveform analysis revealed subtle differences between the PCD or knockdown cells and controls that varied among the cells. The heterogeneous waveform observed between individual cilia in the PCD samples could be related to the maturity of individual cilium. It has been shown that the variation in ciliary waveform is associated with progression of ciliary length and the differentiation of ciliated cells (52).

To characterize the function of CFAP57 in more detail, we utilized the well-studied model organism *Chlamydomonas*. We show here that *CFAP57* encodes a WDR protein that plays a role in the docking/assembly of a subset of IDAs. Although the ciliary axoneme is described as showing nine-fold symmetry, there are many structural asymmetries in the cilium (reviewed in Dutcher, in revision). The generation of waveforms requires the spatial and temporal regulation of dyneins, and this is likely to require structural asymmetries. Multiple approaches have helped to catalog the asymmetric structures and proteins in *Chlamydomonas* cilia. These asymmetries have been identified through analysis of mutant *Chlamydomonas* flagella by electron tomography and proteomics and are beginning to provide a wealth of information to use for understanding how asymmetric and symmetric waveforms are generated and propagated (27, 28, 45). If all of the ciliary dyneins were active at one time, the cilia would be in a rigor state, resulting in no net movement or bending. In order to generate an effective bend, dynein motor function must be tightly controlled both along the length of the cilium and around the circumference of the axoneme across a defined axis.

In elegant cryo-EM tomography studies, Lin and Nicastro unexpectedly found that most dyneins are in an active state confirmation with a smaller population of inactive dyneins (53). The locations of the inactive dyneins are asymmetric and bend-dependent. They propose a switch-inhibition mechanism in which the bend is generated by inhibiting, rather than activating, dyneins on one side of the cilium. The data suggest that the initiation of a bend starts with the inhibition of the inner dynein arms (a, d, and g) on specific doublet microtubules (DMT). DMT 2 to 4 is needed for the asymmetric waveform and DMT 7 to 9 for the symmetrical waveform that promotes a bend by the dyneins on the other side of the axoneme. CFAP57 is likely to play a key role in regulating the waveform.

The electron tomographic analysis of the *Chlamydomonas bop2-1* mutant showed that CFAP57 is required for the assembly of dynein arms on only a subset of the nine doublet microtubules (26). A loss of IDAs on doublets 5, 6, and 8 was observed. In recent work using cryo-EM tomography, there is a loss of a subset of inner dynein arms on doublet microtubules 5-8 with a partial loss on doublets 1 and 9 (Lin et al., (http://dx.doi.org/10.1101/688291). Lin et al. provides data that CFAP57 forms an extended filament that connects to several structures. The proteome analysis of *fap57/bop2* mutants suggests that dynein d (DHC7) and g (DHC2) are reduced in the mutant along with the minor dynein, DHC3, which assembles at the far end of the 96 nm repeat in the proximal region where dynein g is found distally and could be considered the proximal g inner dynein. The loss of these dyneins on only a subset of doublet microtubules is sufficient to affect the swimming velocity and the waveform in *Chlamydomonas.* The CFAP57 filament may act like the CCDC39/CCDC40 ruler that specifies the addresses of the N-DRC and radial spokes (54, 55).

FAP43 and FAP44 are required for the formation of the tether and tether head (T/TH) complex, which is required for the positional stability of the I1/f dynein motor domains, as well as the stable anchoring of CK1 kinase, and proper phosphorylation of the regulatory IC138 subunit in *Chlamydomonas*. Interestingly, T/TH also interacts with the inner dynein arm d and radial spoke 3 (5). In *Tetrahymena,* CFAP57 is placed adjacent to the FAP43/44 complex based on proximity mapping (6). Comparative proteomics analysis of I1/f and FAP43/44 mutants showed that the I1 dynein and the T/TH complex assemble independently of each other (56). The *fap57* mutants assemble the Il/f two-headed dynein complex properly as well (Table 3).

In comparison to *Chlamydomonas*, less is known about the detailed structure of the IDA in human cilia. Based on the high level of conservation between species, it is likely that CFAP57 plays a similar role in human cilia as it does in *Chlamydomonas*. However, because the planar waveform of human cilia is inherently different from the waveforms of *Chlamydomonas*, the exact positioning and regulation of IDA activity is also likely different. Additional studies using advanced techniques to culture and study human respiratory cells are needed to address these questions.

In summary, our results show that a genetic variant in *CFAP57* reduces CBF and alters waveform, likely by affecting the assembly of a subset of IDA, resulting in PCD. This is the first reported example of PCD caused by mutation of a protein that apparently affects only a subset of the IDAs, expanding our understanding of how dynein arms are positioned during cilia assembly. These findings demonstrate the usefulness of the PSAP algorithm and set a precedent to consider it in the evaluation of other cases of PCD with no obvious structural defects. Identifying the genetic basis of PCD and the functional defects is an important step toward developing personalized treatments for this rare disease.

## Methods

### Human subjects

The individuals included in this study provided informed consents and all protocols involving human studies were approved by the University of North Carolina Medical School Institutional Review Board.

### Human genetic analysis

A cohort of 99 PCD patients with no obvious EM phenotype was assembled for exome sequencing. Exome libraries were prepared using an IDT capture reagent. Genetic variants were discovered and genotyped using a validated analysis pipeline at the Washington University McDonnell Genome Institute as previously described (31). Each case was analyzed using the population sampling probability (PSAP) framework, a published statistical method for identifying pathogenic mutations from n=1 cases of rare disease (24). Segregation analysis was performed on the available DNA from family members (family UNC-1095). The primers used are listed in Table S6.

### Airway epithelial cell cultures

***Human nasal epithelial (HNE) cells*** from the PCD subject (proband 2-II) and controls were obtained as described (7). The nasal cells were expanded as conditionally reprogrammed cells (CRC) (8) and cultured as previously described (9).

***Human tracheobronchial epithelial cells (hTEC)*** were obtained from non-smoking donors lacking respiratory pathologies provided by the Cystic Fibrosis Center Tissue Procurement and Cell Culture Core (59), or were isolated from surgical excess of tracheobronchial segments of lungs donated for transplantation as previously described (11). These unidentified cells are exempt from regulation by HHS regulation 45 CFR Part 46. hTEC cells were expanded *in vitro* and allowed to differentiate using air-liquid interface (ALI) conditions on supported membranes (Transwell, Corning Inc., Corning, NY), as previously described (11, 32, 59). These protocols have been approved by the University of North Carolina (UNC) Medical School Institutional Review Board.

#### shRNA for silencing of *CFAP57*

shRNA sequences were obtained from Sigma-Aldrich in pLK0.1 lentivirus vector, and were chosen from the Broad institute RNAi library. Target sequences evaluated included (Sequence 1: CCTGGAACTATTTAAGGAATA, Sequence 2: CAGAAAGTAATGGCCATTGTT, sequence 3: CCCTGGAACTATTTAAGGAAT, sequence 4: CCTTCCATTCACCCTTCTCAT, sequence 5: GCCTAGGAAATCATCAGAGA).

#### Analysis of *CFAP57* expression

RNA was isolated from cells using an RNeasy Mini Kit (Qiagen) and RT-PCR was performed using specific primers (Table S6) as previously described (22). For qRT-PCR, RNA was reverse-transcribed using an Applied Bioscience High-Capacity Reverse Transcription Kit (Thermo Fisher Scientific). Gene expression was detected by RT-PCR using gene specific primers, (FW:AAAGCAGAACTGTTTGGCGG, RV:TTGGGACTGATGGACAAGGC) and a SYBR green nucleic acid-labeling SYBR FAST kit (Kapa Biosystems) in a Lightcycler 480 (Roche). Fold change was calculated using the delta–delta cycle threshold [ΔΔC(t)] analysis method and OAZ1 as an internal control.

For RNAseq analysis, hTEC cells were cultured using ALI conditions, and sampled at ALI day 7, 14, and 21. Three samples from each time point were used for analysis. RNA was extracted using a Qiagen RNeasy kit. RNA library preparation and analysis were performed by the Washington University Genome Center (GTAC), using Clontech-SMARTer RNAseq kit, and sequenced using an Illumina Hiseq3000 sequencer, for a total depth reads of at least 40 million reads per sample.

For *Chlamydomonas* RNA isolation, cells from two R medium agar plates grown for 5 days were resuspended in 40 ml nitrogen-free medium (M-N/5) for 2 h at room temperature to allow flagellar assembly. The cells were then collected, and RNA extraction was performed with the RNeasy Mini Kit (Qiagen) according to the manufacturer’s recommendation. Two micrograms of RNA was used in a reverse transcription reaction with SuperScript III (Invitrogen) with random primers as previously described (60).

#### Epithelial cell immunofluorescence

Airway cells were fixed and immunostained as previously described (32, 35, 61). The binding of primary antibodies was detected using Alexa Fluor-488, Alexa Fluor-647 (Life Technologies), indocarbocyanine (CY3) or Rhodamine Red-X (RRX) conjugated secondary antibody (Jackson ImmunoResearch Laboratories, West Grove, PA). The DNA was stained using 4’, 6-diamidino-2-phenylindole (DAPI, Vector Laboratories, Burlingame, CA, USA) or with Hoechst 33342 FluoroPure™ (Life Technologies). Slides were mounted using ProLong Diamond antifade mountant (Thermo Fisher). Images were acquired using a Zeiss -710 microscopy system or a Leica epifluorescent microscope (LAS X, Leica, Buffalo Grove, IL) Images were processed and the fluorescence intensity analyzed using FIJI (10) as previously described (11). Brightness and contrast were adjusted globally using Photoshop (Adobe Systems, San Jose, CA). Isotype matched control antibodies had no detectable staining under the conditions used. The antibodies used are listed in Table S5.

#### Ciliary isolation, protein extraction and Immunoblot

Ciliary isolation and protein extraction was performed as previously described (35). Protein extraction from tissue or cells and immunoblot analysis was performed as previously described (35, 61).

#### Ciliary beat frequency and waveform analysis

The ciliary beat frequency (CBF) in HNE cultures (n=6) was measured as previously described (22, 63). In hTEC cultures, CBF was measured in at least 5 fields obtained from each preparation. The cultures were maintained at 37 °C using a temperature controller and a stage heater block. High-speed videos (120 frames/s) were recorded and processed with the Sisson-Ammons Video Analysis system (SAVA, Amons Engineering, Mt Morris, MI) as described (11, 64). To analyze the ciliary waveform, cells were lifted off the supportive membranes and imaged directly on slides. High resolution videos of 6 cells in 3 different cultures were recorded as previously described (22). The videos were analyzed by an experienced scientist blinded to the genotype of the cells. Videos were replayed in slow motion. The ciliary length was measured and the waveform of the front and back cilia in 4 ciliated cells was traced manually. Deviation from linearity was determined by measuring the furthest ciliary displacement from the linear axis as defined by the end-effective and end-recovery position, as previously described (65).

#### *Chlamydomonas* axoneme isolation and mass spectroscopy

*Chlamydomonas* cilia were isolated using the dibucaine method (66). Cilia were demembranated with the addition of 1% Nonidet P-40 in HMDS-EGTA (10 mM HEPES, 5 mM MgSO_4_, 1 mM DTT, 4% Sucrose, 0.5 mM EGTA, pH7.4) buffer and separated from the membrane and matrix fractions by centrifugation. Isolated cilia were resuspended in HMDEK (30 mM HEPES, 5 mM MgSO_4_ 1 mM DTT, 0.5 mM EGTA, 25 mM KCl, pH 7.45) buffer. 100 micrograms of proteins from each sample were digested by trypsin and given isobaric tags before undergoing mass spectrometry (Donald Danforth Plant Science Center, St. Louis, MO).

#### Preparation of *Chlamydomonas* for video microscopy

*Chlamydomonas reinhardtii* cells were grown as previously described (67). Cells were grown on agar plate for 48 hours in Sager and Granick rich liquid medium (68) at 25°C in constant light. Prior to recording, cells were suspended for 3 hours in a medium lacking nitrogen adapted from Medium I of Sager and Granick (68) to promote gametogenesis. Cells were directly recording under the microscope.

All bright field microscopy was carried in a climate-control room maintained at 21°C. For each recording, 10 μL from liquid cells culture (after gametogenesis) were pipetted onto a slide and a cover slip (18 x 18 mm) was placed for recording under a Zeiss Axiophot (Carl Zeiss AG, Oberkochen, Germany) with a 40x Plan-Neofluar objective lens for swimming analysis and with 100x Neofluar oil-immersion objective lens for waveform analysis. Videos were recording using a Phantom Miro eX2 camera and Phantom Camera Control Application 2.6 (Vision Research©, Inc, Wayne, NJ, USA). Videos were captured at 2000 frames per second with 320 x 240 resolution and an exposure time of 200 μs. Around 7000 frames with 3500 frames before the trigger and 3500 frames after the trigger were captured. Frames displaying a characteristic beating/swimming were extracted and saved under uncompressed AVI format at 15 frames per second.

#### *Chlamydomonas* swimming analysis

Each video were analyzed using ImageJ (69) to create a binary file only displaying the cells. Each pixel had a spatial resolution of 310 x 310 nm and the temporal resolution between 2 consecutive time points was 1 ms. Cells were tracked using the 2D/3D single-particle tracking tool of the MosaicSuite for ImageJ and Fiji (©MOSAIC Group, MPI-CBG, Dresden). Custom-made program written in Matlab™ R2016a (The Mathworks, Natick, MA, USA) was then used to compute the cells velocity as well as the ciliary beat frequency extracted by Fast-Fourier-Transform of the trajectory.

#### *Chlamydomonas* kinematic analysis of cilium

The uniciliate mutant strain *uni1-2* and the double mutants *fap57-050; uni1* and *fap57-706; uni1* were generated from meiotic crosses, as described by Dutcher (70). The single mutant *uni1* is considered the wild-type reference as previously published (43). Videos were analyzed using a custom-made program written in Matlab™ R2016a (The Mathworks, Natick, MA, USA) previously published (44). From each video, a sequence of 200 consecutive frames was stored in a 3D matrix of pixel intensity values. Each pixel had a spatial resolution of 169 x 169 nm and the temporal resolution between 2 consecutive time points was 0.5 ms. Components of the forces exerted by cilia on the fluid are directly calculated from the Cartesian coordinates using parameters previously reported (44).

#### Statistical analyses

Group variation is described as mean ± standard error (SEM). Statistical comparisons between groups were made using 1-way analysis of variance (ANOVA) with Tukey post-hoc analysis. Individual group differences were determined using a 2-tailed Student’s t-test. A p value of 0.05 was considered to represent a significant difference. Non-parametric data are shown as the median and the 25^th^ and 75^th^ intraquartile ranges. Data were analyzed using Prism (GraphPad, La Jolla, CA).

### TMT mass spectroscopy

#### Protein extraction and digestion

The samples were lysed in lysis buffer (8M Urea in 50mM HEPES pH 8.0) and sonicated briefly. Samples were reduced with 10 mM TCEP and alkylated with 25 mM iodoacetamide. Protein concentrations were determined by BCA protein assay (Thermo Scientific). Fifty μg of protein was taken from each sample for digestion. Sequencing grade protease Lys-C was added with 1:250 ratio and sample was digested overnight at room temperature with mixing. A second digestion was performed by diluting the sample with 50 mM HEPES to lower the urea concentration to 1M and trypsin was added with 1:100 ratio for a further 12 hour digestion at room temperature with mixing. Digests were acidified with formic acid and subjected to Oasis HLB solid phase extraction column (Waters).

#### Tandem Mass Tag (TMT) Labeling

Digested peptides were labeled according to the TMT 10plex reagent kit instructions. Briefly, TMT regents were brought to room temperature and dissolved in anhydrous acetonitrile. Peptides were labeled by the addition of each label to its respective digested sample. Labeling reactions were incubated without shaking for 1 h at room temperature. Reactions were terminated with the addition of hydroxylamine. Subsequent labeled digests were combined into a new 2 mL microfuge tube, acidified with formic acid, subjected to Sep-Pak C18 solid phase extraction and dried down.

#### High pH Reverse Phase Fractionation

The dried peptide mixture was dissolved in 110 μL of mobile phase A (10 mM ammonium formate, pH 9.0). 100 μL of the sample was injected onto a 2.1 x 150 mm XSelect CSH C18 column (Waters) equilibrated with 3% mobile phase B (10 mM ammonium formate, 90% ACN). Peptides were separated using a gradient (71) to at a flow rate of 0.2 mL/min. 60 peptide fractions were collected corresponding to 1 min each. 10 pooled samples were generated by concatenation in which every 10th fraction (ie: 1, 11, 21, 31, 41, 51; six fractions total) was combined. The 10 pooled samples were acidified and dried down prior to LC-MS analysis. LC-MS Analysis Each fraction was resuspended in 50 μL 1% acetonitrile/1% formic acid. 5 μL was analyzed by LC-MS (HCD for MS/MS) with a Dionex RSLCnano HPLC coupled to a Velos Pro OrbiTrap mass spectrometer (Thermo Scientific) using a 2h gradient. Peptides were resolved using 75 μm x 25 cm PepMap C18 column (Thermo Scientific).

#### Data Analysis

All MS/MS samples were analyzed using Proteome Discoverer 2.1 (Thermo Scientific). The Sequest HT search engine in the Proteome Discover was set to search *Chlamydomonas reinhardtii* database (Uniprot.org). The digestion enzyme was set as trypsin. The HCD MS/MS spectra were searched with a fragment ion mass tolerance of 0.02 Da and a parent ion tolerance of 20 ppm. Oxidation of methionine was specified as a variable modification, while carbamidomethyl of cysteine and TMT labeling was designated at lysine residues or peptide N-termini were specified in Proteome Discoverer as static modifications.

MS/MS based peptide and protein identifications and quantification results was initially generated in Proteome Discover 2.1 and later uploaded to Scaffold (version Scaffold_4.8.2 Proteome Software Inc., Portland, OR) for final TMT quantification and data visualization. Normalized and scaled protein/peptide abundance ratios were calculated against the abundance value of the four wild-type controls. Peptide identifications were accepted if they could be established at greater than 80.0% probability by the Peptide Prophet algorithm (12) with Scaffold delta-mass correction. Protein identifications were accepted if they could be established at greater than 99.0% probability and contained at least 2 identified peptides. Protein probabilities were assigned by the Protein Prophet algorithm (73). Proteins that contained similar peptides and could not be differentiated based on MS/MS analysis alone were grouped to satisfy the principles of parsimony. Proteins sharing significant peptide evidence were grouped into clusters.

## Supporting information

Legends for movies

## Acknowledgments

We are grateful to the patient and the family members, Michele Manion (founder of the US PCD Foundation), and the US PCD foundation. We thank investigators and the coordinators of the Genetic Disorders of Mucociliary Clearance Consortium that is part of the Rare Disease Clinical Research Network; Whitney Wolf, Kimberly Burns from UNC for technical assistance and Hong Dang from UNC for bioinformatics assistance. Drs. Shrikant Mane and Francesc Lopez-Giraldez, and Ms. Weilai Dong from Yale Center for Mendelian Genomics for providing whole exome sequencing and bioinformatics support. We thank the McDonnell Genome Institute for their expertise in whole exome sequencing.

Funding support for research was provided to S.K.D. by US NIH-NHLBI grant R01HL128370; to M.R.K., L.A.D, M.W.L., and M.A.Z by The Genetic Disorders of Mucociliary Clearance (U54HL096458), a part of the NCATS Rare Diseases Clinical Research Network (RDCRN). RDCRN is an initiative of the Office of Rare Diseases Research (ORDR), NCATS, funded through a collaboration between NCATS and NHLBI. To A.H. by the American Thoracic Society Foundation/Primary Ciliary Dyskinesia Foundation/Kovler Family Foundation joint grant; to M.R.K., L.E.O., and M.A.Z. by NIH-NHLBI grant R01HL071798, to L.E.O., M.R.K., and M.A.Z. by NIH/NHLBI R01 HL117836, to University of North Carolina at Chapel Hill by grant UL1 TR000083 from the NCATS at NIH. D.F.C was funded by NIH/NICHD R01HD078641 and NIH/NIMH R01MH101810. The Yale Center for Mendelian Genomics (UM1HG006504) is funded by the NHGRI. The GSP Coordinating Center (U24 HG008956) contributed to cross-program scientific initiatives and provided logistical and general study coordination. The contents are solely the responsibility of the authors and do not necessarily represent the official views of the National Institute of Health.

## References

1. Ferkol TW & Leigh MW (2012) Ciliopathies: the central role of cilia in a spectrum of pediatric disorders. The Journal of pediatrics 160(3):366–371.

2. Zariwala MA, Knowles MR, & Leigh MW (2007 [Updated 2015]) Primary Ciliary Dyskinesia. GeneReviews(R) [Internet], eds Pagon RA, Adam MP, Ardinger HH, Wallace SE, Amemiya A, Bean LJH, Bird TD, Fong CT, Mefford HC, Smith RJH, et al. (University of Washington, Seattle 1993-2016. Available from: http://www.ncbi.nlm.nih.gov/books/NBK1122/, Seattle (WA)).

3. Shapiro AJ, et al. (2016) Diagnosis, monitoring, and treatment of primary ciliary dyskinesia: PCD foundation consensus recommendations based on state of the art review. Pediatric pulmonology 51(2):115–132.

4. Knowles MR, et al. (2012) Mutations of DNAH11 in patients with primary ciliary dyskinesia with normal ciliary ultrastructure. Thorax 67(5):433–441.

5. Knowles MR, et al. (2013) Exome sequencing identifies mutations in CCDC114 as a cause of primary ciliary dyskinesia. American journal of human genetics 92(1):99–106.

6. Hjeij R, et al. (2014) CCDC151 mutations cause primary ciliary dyskinesia by disruption of the outer dynein arm docking complex formation. American journal of human genetics 95(3):257–274.

7. Alsaadi MM, et al. (2014) Nonsense mutation in coiled-coil domain containing 151 gene (CCDC151) causes primary ciliary dyskinesia. Human mutation 35(12):1446–1448.

8. Mitchison HM, et al. (2012) Mutations in axonemal dynein assembly factor DNAAF3 cause primary ciliary dyskinesia. Nature genetics 44(4):381–389, S381-382.

9. Panizzi JR, et al. (2012) CCDC103 mutations cause primary ciliary dyskinesia by disrupting assembly of ciliary dynein arms. Nature genetics 44(6):714–719.

10. Tarkar A, et al. (2013) DYX1C1 is required for axonemal dynein assembly and ciliary motility. Nature genetics 45(9):995–1003.

11. Horani A, et al. (2012) Whole-Exome Capture and Sequencing Identifies HEATR2 Mutation as a Cause of Primary Ciliary Dyskinesia. American journal of human genetics 91(4):685–693.

12. Horani A, et al. (2013) LRRC6 mutation causes primary ciliary dyskinesia with dynein arm defects. PloS one 8(3):e59436.

13. Horani A, et al. (2018) Establishment of the early cilia preassembly protein complex during motile ciliogenesis. Proceedings of the National Academy of Sciences of the United States of America 115(6):E1221–E1228.

14. Knowles Michael R, et al. (2013) Mutations in SPAG1 Cause Primary Ciliary Dyskinesia Associated with Defective Outer and Inner Dynein Arms. American journal of human genetics 93(4):711–720.

15. Wallmeier J, et al. (2014) Mutations in CCNO result in congenital mucociliary clearance disorder with reduced generation of multiple motile cilia. Nature genetics 46(6):646–651.

16. Boon M, et al. (2014) MCIDAS mutations result in a mucociliary clearance disorder with reduced generation of multiple motile cilia. Nature communications 5:4418.

17. Olbrich H, et al. (2015) Loss-of-Function GAS8 Mutations Cause Primary Ciliary Dyskinesia and Disrupt the Nexin-Dynein Regulatory Complex. American journal of human genetics 97(4):546–554.

18. Edelbusch C, et al. (2017) Mutation of serine/threonine protein kinase 36 (STK36) causes primary ciliary dyskinesia with a central pair defect. Human mutation 38(8):964–969.

19. Wallmeier J, et al. (2016) TTC25 Deficiency Results in Defects of the Outer Dynein Arm Docking Machinery and Primary Ciliary Dyskinesia with Left-Right Body Asymmetry Randomization. The American Journal of Human Genetics 99(2):460–469.

20. Olcese C, et al. (2017) X-linked primary ciliary dyskinesia due to mutations in the cytoplasmic axonemal dynein assembly factor PIH1D3. 8:14279.

21. Paff T, et al. (2017) Mutations in PIH1D3 Cause X-Linked Primary Ciliary Dyskinesia with Outer and Inner Dynein Arm Defects. The American Journal of Human Genetics 100(1):160–168.

22. Bustamante-Marin XM, et al. (2019) Lack of GAS2L2 Causes PCD by Impairing Cilia Orientation and Mucociliary Clearance. The American Journal of Human Genetics. 104(2):229–245.

23. El Khouri E, et al. (2016) Mutations in DNAJB13, Encoding an HSP40 Family Member, Cause Primary Ciliary Dyskinesia and Male Infertility. The American Journal of Human Genetics 99(2):489–500.

24. Wilfert AB, et al. (2016) Genome-wide significance testing of variation from single case exomes. Nature genetics 48(12):1455–1461.

25. Dutcher SK, Gibbons W, & Inwood WB (1988) A genetic analysis of suppressors of the PF10 mutation in Chlamydomonas reinhardtii. Genetics 120(4):965–976.

26. King SJ, Inwood WB, O’Toole ET, Power J, & Dutcher SK (1994) The bop2-1 mutation reveals radial asymmetry in the inner dynein arm region of Chlamydomonas reinhardtii. The Journal of cell biology 126(5):1255–1266.

27. Bui KH, Yagi T, Yamamoto R, Kamiya R, & Ishikawa T (2012) Polarity and asymmetry in the arrangement of dynein and related structures in the Chlamydomonas axoneme. The Journal of cell biology 198(5):913–925.

28. Heuser T, et al. (2012) Cryoelectron tomography reveals doublet-specific structures and unique interactions in the I1 dynein. Proceedings of the National Academy of Sciences of the United States of America 109(30):E2067–2076.

29. Yagi T, Uematsu K, Liu Z, & Kamiya R (2009) Identification of dyneins that localize exclusively to the proximal portion of Chlamydomonas flagella. Journal of cell science 122(Pt 9):1306–1314.

30. Leigh MW, et al. (2013) Standardizing Nasal Nitric Oxide Measurement as a Test for Primary Ciliary Dyskinesia. Annals of the American Thoracic Society 10(6):574–581.

31. Griffith M, et al. (2015) Optimizing cancer genome sequencing and analysis. Cell Syst 1(3):210–223.

32. You Y, Richer EJ, Huang T, & Brody SL (2002) Growth and differentiation of mouse tracheal epithelial cells: selection of a proliferative population. Am J Physiol Lung Cell Mol Physiol 283(6):L1315–1321.

33. Horani A, Nath A, Wasserman MG, Huang T, & Brody SL (2013) Rho-associated protein kinase inhibition enhances airway epithelial Basal-cell proliferation and lentivirus transduction. Am J Respir Cell Mol Biol 49(3):341–347.

34. Pazour GJ, Agrin N, Leszyk J, & Witman GB (2005) Proteomic analysis of a eukaryotic cilium. The Journal of cell biology 170(1):103–113.

35. Blackburn K, Bustamante-Marin X, Yin W, Goshe MB, & Ostrowski LE (2017) Quantitative Proteomic Analysis of Human Airway Cilia Identifies Previously Uncharacterized Proteins of High Abundance. J Proteome Res 16(4):1579–1592.

36. Gentzsch M, et al. (2017) Pharmacological Rescue of Conditionally Reprogrammed Cystic Fibrosis Bronchial Epithelial Cells. American journal of respiratory cell and molecular biology 56(5):568–574.

37. Feng Y, et al. (A multifunctional lentiviral-based gene knockdown with concurrent rescue that controls for off-target effects of RNAi. Genomics Proteomics Bioinformatics 8(4):238–245.

38. Horani A, et al. (2012) Whole-Exome Capture and Sequencing Identifies HEATR2 Mutation as a Cause of Primary Ciliary Dyskinesia. American journal of human genetics 91(4):685–693.

39. Yachdav G, et al. (2014) PredictProtein--an open resource for online prediction of protein structural and functional features. Nucleic acids research 42(Web Server issue):W337–343.

40. Dutcher SK (2000) Chlamydomonas reinhardtii: biological rationale for genomics. The Journal of eukaryotic microbiology 47(4):340–349.

41. Li X, et al. (2016) An Indexed, Mapped Mutant Library Enables Reverse Genetics Studies of Biological Processes in Chlamydomonas reinhardtii. The Plant cell 28(2):367–387.

42. Zhang R, et al. (2014) High-Throughput Genotyping of Green Algal Mutants Reveals Random Distribution of Mutagenic Insertion Sites and Endonucleolytic Cleavage of Transforming DNA. The Plant cell 26 (4):1398–1409.

43. Bayly PV, Lewis BL, Kemp PS, Pless RB, & Dutcher SK (2010) Efficient spatiotemporal analysis of the flagellar waveform of Chlamydomonas reinhardtii. Cytoskeleton 67(1):56–69.

44. Bottier M, Thomas KA, Dutcher SK, & Bayly PV (2019) How Does Cilium Length Affect Beating? Biophysical Journal 116(7):1292–1304.

45. Lin H, Guo S, & Dutcher SK (2018) RPGRIP1L helps to establish the ciliary gate for entry of proteins. Journal of cell science 131(20): jcs220905.

46. Bower R, et al. (2013) The N-DRC forms a conserved biochemical complex that maintains outer doublet alignment and limits microtubule sliding in motile axonemes. Mol Biol Cell 24(8):1134–1152.

47. Awata J, et al. (2014) NPHP4 controls ciliary trafficking of membrane proteins and large soluble proteins at the transition zone. Journal of cell science 127(Pt 21):4714–4727.

48. Consortium GT (2015) Human genomics. The Genotype-Tissue Expression (GTEx) pilot analysis: multitissue gene regulation in humans. Science 348(6235):648–660.

49. Horani A, Ferkol TW, Dutcher SK, & Brody SL (2016) Genetics and biology of primary ciliary dyskinesia. Paediatr Respir Rev 18:18–24.

50. Kasak L, et al. (2018) Bi-allelic Recessive Loss-of-Function Variants in FANCM Cause Non-obstructive Azoospermia. Am J Hum Genet 103(2):200–212.

51. Lin J & Nicastro D (2018) Asymmetric distribution and spatial switching of dynein activity generates ciliary motility. Science (New York, N.Y.) 360(6387):eaar1968.

52. Oltean A, Schaffer AJ, Bayly PV, & Brody SL (2018) Quantifying Ciliary Dynamics during Assembly Reveals Stepwise Waveform Maturation in Airway Cells. American journal of respiratory cell and molecular biology 59(4):511–522.

53. Lin J & Nicastro D (2018) Asymmetric distribution and spatial switching of dynein activity generates ciliary motility. Science 360(6387).

54. Oda T, Yanagisawa H, Kamiya R, & Kikkawa M (2014) A molecular ruler determines the repeat length in eukaryotic cilia and flagella. Science 346(6211):857–860.

55. Lin H, et al. (2015) A NIMA-Related Kinase Suppresses the Flagellar Instability Associated with the Loss of Multiple Axonemal Structures. PLoS Genet 11(9):e1005508.

56. Fu G, et al. (2018) The I1 dynein-associated tether and tether head complex is a conserved regulator of ciliary motility. Mol Biol Cell 29(9):1048–1059.

57. Urbanska P, et al. (2018) Ciliary proteins Fap43 and Fap44 interact with each other and are essential for proper cilia and flagella beating. Cellular and molecular life sciences : CMLS 75(24):4479–4493.

58. Muller L, Brighton LE, Carson JL, Fischer WA, 2nd, & Jaspers I (2013) Culturing of human nasal epithelial cells at the air liquid interface. J Vis Exp (80). doi: 10.3791/50646.

59. Fulcher ML & Randell SH (2013) Human nasal and tracheo-bronchial respiratory epithelial cell culture. Methods Mol Biol 945:109–121.

60. Lin H, Kwan AL, & Dutcher SK (2010) Synthesizing and salvaging NAD: lessons learned from Chlamydomonas reinhardtii. PLoS Genet 6(9):e1001105.

61. Pan J, You Y, Huang T, & Brody SL (2007) RhoA-mediated apical actin enrichment is required for ciliogenesis and promoted by Foxj1. Journal of cell science 120(Pt 11):1868–1876.

62. Schindelin J, et al. (2012) Fiji: an open-source platform for biological-image analysis. Nature methods 9:676.

63. Sears PR, Yin WN, & Ostrowski LE (2015) Continuous mucociliary transport by primary human airway epithelial cells in vitro. American journal of physiology. Lung cellular and molecular physiology 309(2):L99–108.

64. Sisson JH, Stoner JA, Ammons BA, & Wyatt TA (2003) All-digital image capture and whole-field analysis of ciliary beat frequency. J Microsc 211(Pt 2):103–111.

65. Knowles MR, et al. (2014) Mutations in RSPH1 Cause Primary Ciliary Dyskinesia with a Unique Clinical and Ciliary Phenotype. American journal of respiratory and critical care medicine 189(6):707–717.

66. Witman GB (1986) Isolation of Chlamydomonas flagella and flagellar axonemes. Methods in enzymology 134:280–290.

67. Holmes JA & Dutcher SK (1989) Cellular asymmetry in Chlamydomonas reinhardtii. Journal of cell science 94 (Pt 2):273–285.

68. Lux FG, 3rd & Dutcher SK (1991) Genetic interactions at the FLA10 locus: suppressors and synthetic phenotypes that affect the cell cycle and flagellar function in Chlamydomonas reinhardtii. Genetics 128(3):549–561.

69. Schneider CA, Rasband WS, & Eliceiri KW (2012) NIH Image to ImageJ: 25 years of image analysis. Nature methods 9(7):671–675.

70. Dutcher SK (1995) Flagellar assembly in two hundred and fifty easy-to-follow steps. Trends in genetics : TIG 11(10):398–404.

71. Batth TS, Francavilla C, & Olsen JV (2014) Off-line high-pH reversed-phase fractionation for in-depth phosphoproteomics. J Proteome Res 13(12):6176–6186.

72. Keller A, Nesvizhskii AI, Kolker E, & Aebersold R (2002) Empirical statistical model to estimate the accuracy of peptide identifications made by MS/MS and database search. Analytical chemistry 74(20):5383–5392.

73. Nesvizhskii AI, Keller A, Kolker E, & Aebersold R (2003) A statistical model for identifying proteins by tandem mass spectrometry. Analytical chemistry 75(17):4646–4658.

## References

1. Kagami O & Kamiya R (1995) Separation of dynein species by high-pressure liquid chromatography. Methods Cell Biol 47:487–489.

2. Leigh MW, et al. (2013) Standardizing Nasal Nitric Oxide Measurement as a Test for Primary Ciliary Dyskinesia. Annals of the American Thoracic Society 10(6):574–581.

3. Bayly PV, Lewis BL, Kemp PS, Pless RB, & Dutcher SK (2010) Efficient spatiotemporal analysis of the flagellar waveform of Chlamydomonas reinhardtii. Cytoskeleton 67(1):56–69.

4. Lin J & Nicastro D (2018) Asymmetric distribution and spatial switching of dynein activity generates ciliary motility. Science 360(6387).

5. Fu G, et al. (2018) The I1 dynein-associated tether and tether head complex is a conserved regulator of ciliary motility. Mol Biol Cell 29(9):1048–1059.

6. Urbanska P, et al. (2018) Ciliary proteins Fap43 and Fap44 interact with each other and are essential for proper cilia and flagella beating. Cellular and molecular life sciences : CMLS 75(24):4479–4493.

7. Muller L, Brighton LE, Carson JL, Fischer WA, 2nd, & Jaspers I (2013) Culturing of human nasal epithelial cells at the air liquid interface. J Vis Exp (80).

8. Gentzsch M, et al. (2017) Pharmacological Rescue of Conditionally Reprogrammed Cystic Fibrosis Bronchial Epithelial Cells. American journal of respiratory cell and molecular biology 56(5):568–574.

9. Bustamante-Marin XM, et al. (2019) Lack of GAS2L2 Causes PCD by Impairing Cilia Orientation and Mucociliary Clearance. The American Journal of Human Genetics.

10. Schindelin J, et al. (2012) Fiji: an open-source platform for biological-image analysis. Nature methods 9:676.

11. Blackburn K, Bustamante-Marin X, Yin W, Goshe MB, & Ostrowski LE (2017) Quantitative Proteomic Analysis of Human Airway Cilia Identifies Previously Uncharacterized Proteins of High Abundance. J Proteome Res 16(4):1579–1592.

12. Keller A, Nesvizhskii AI, Kolker E, & Aebersold R (2002) Empirical statistical model to estimate the accuracy of peptide identifications made by MS/MS and database search. Analytical chemistry 74(20):5383–5392.

