## Supplemental tables and figures for "Mutation of *CFAP57* causes primary ciliary dyskinesia by disrupting the asymmetric targeting of a subset of ciliary inner dynein arms"

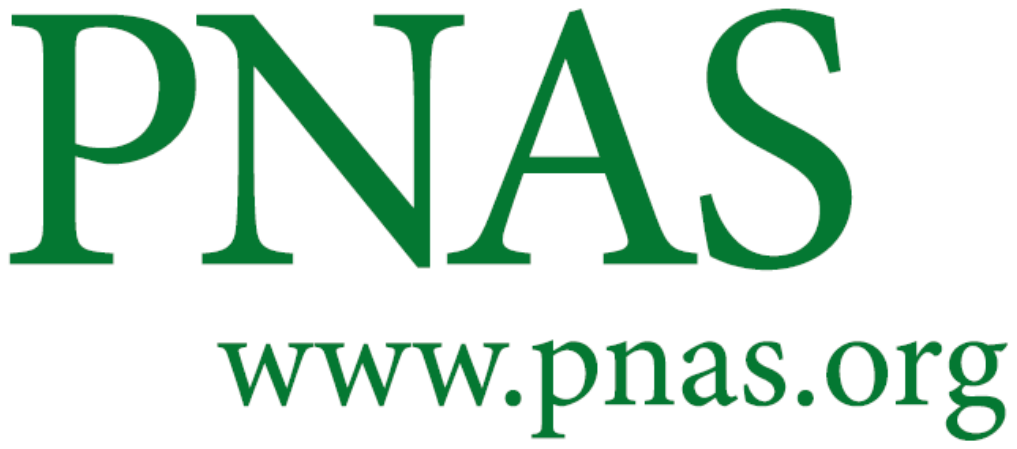

Supplementary Information for

**Mutation of *CFAP57* causes primary ciliary dyskinesia by disrupting the asymmetric targeting of a subset of ciliary inner dynein arm**

Ximena M. Bustamante-Marin, Amjad Horani, Mihaela Stoyanova, Wu-Lin Charng, Mathieu Bottier, Patrick R.. Sears, Leigh Anne Daniels, Hailey Bowe, Donald F. Conrad, Michael R. Knowles, Lawrence E. Ostrowski, Maimoona A. Zariwala and Susan K. Dutcher

Susan K. Dutcher

**This PDF file includes**

Figures S1 to S6

Tables S1 to S6

Legends for Movies S1 to S6

SI References

**Other supplementary materials for this manuscript include the following:**

Movies S1 to S6

**Fig. S1.CFAP57 in non-cultured cells.**

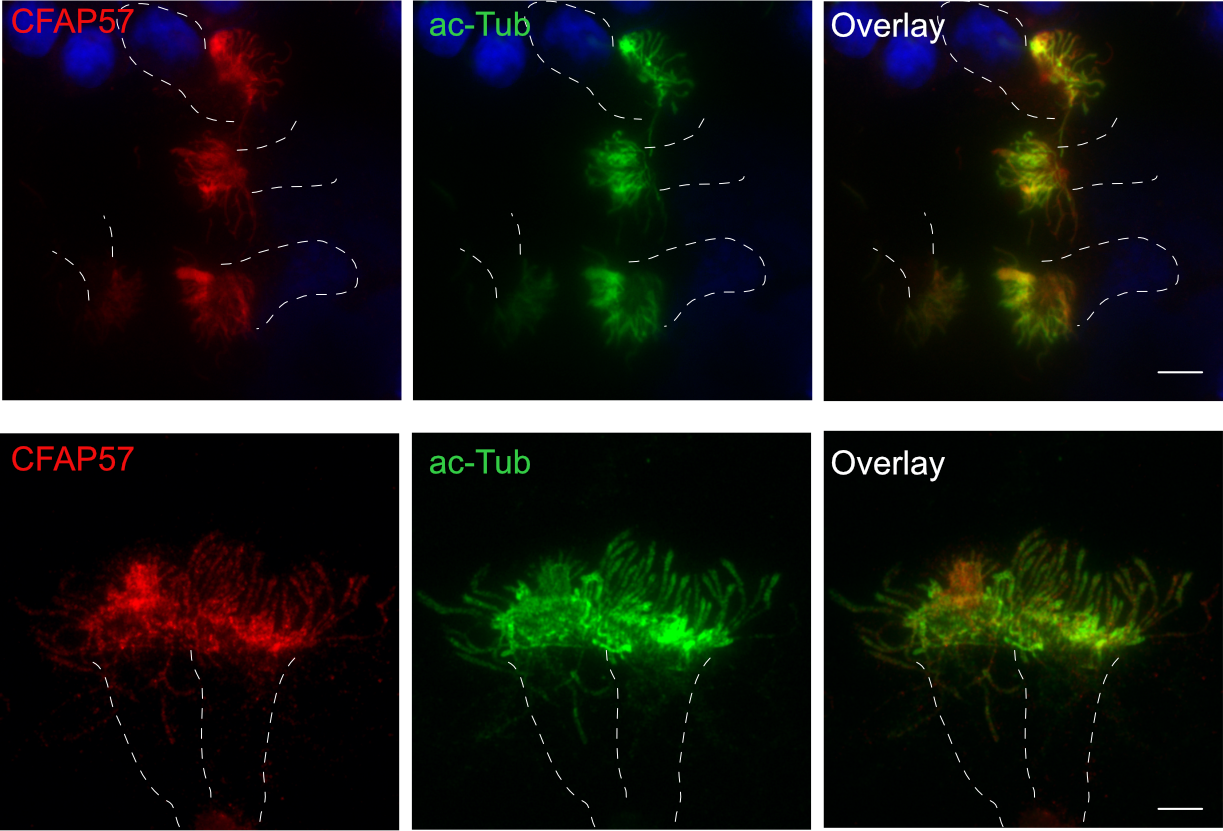

Images showing examples of CFAP57 in non-cultured normal tracheal ciliated cells. Cells were scrapped of fresh human trachea, immunostained with acetylated alpha-tubulin and CFAP57 as detailed in the Materials and Methods.

**Figure** **S2. Immunofluorescence analysis of axonemal components in PCD 2-II cells.**

The staining of isolated ciliated cells from control and PCD 2-II cultures showed normal distribution of (A) DNAH5 (n=3) and (B) RSPH1 (n=3) in different regions of interest (ROI: proximal, middle, and tip) of the ciliary axoneme. The corresponding graph shows the corrected total

cilia fluorescence (CTCF) in each ROI. C)Analysis of the fluorescence intensity for DNALI1 in isolated human nasal ciliated cells obtained from a control (n=5) and the PCD 2-II subject (n=13). These data suggest a reduction of the localization of DNALI1 at the tip of the cilia (p = 0.03

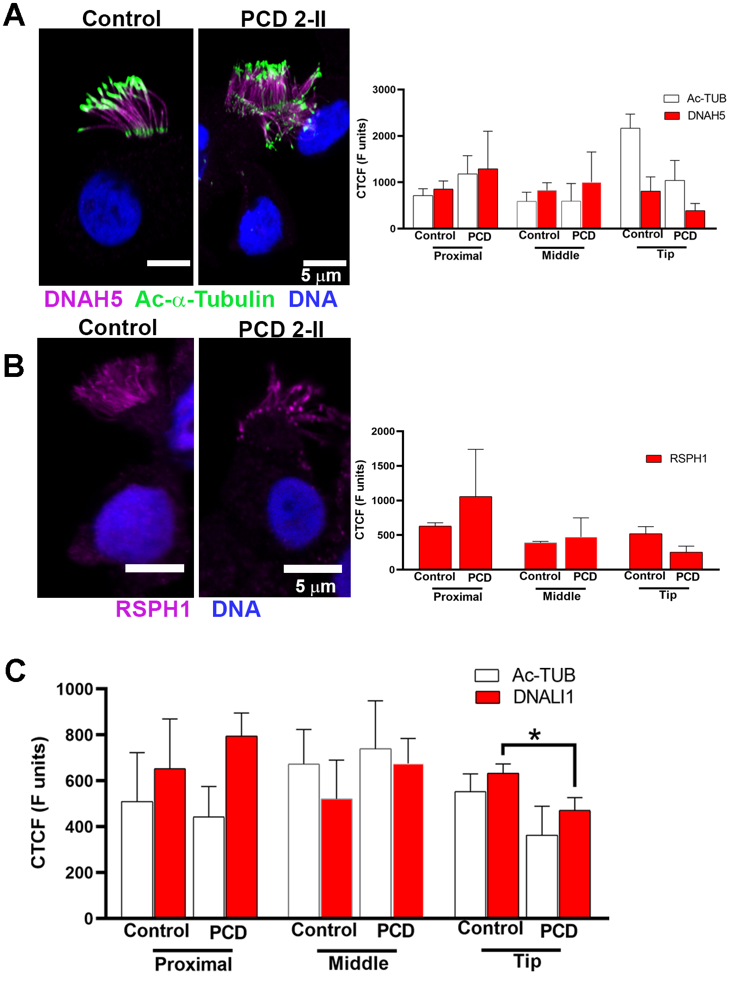

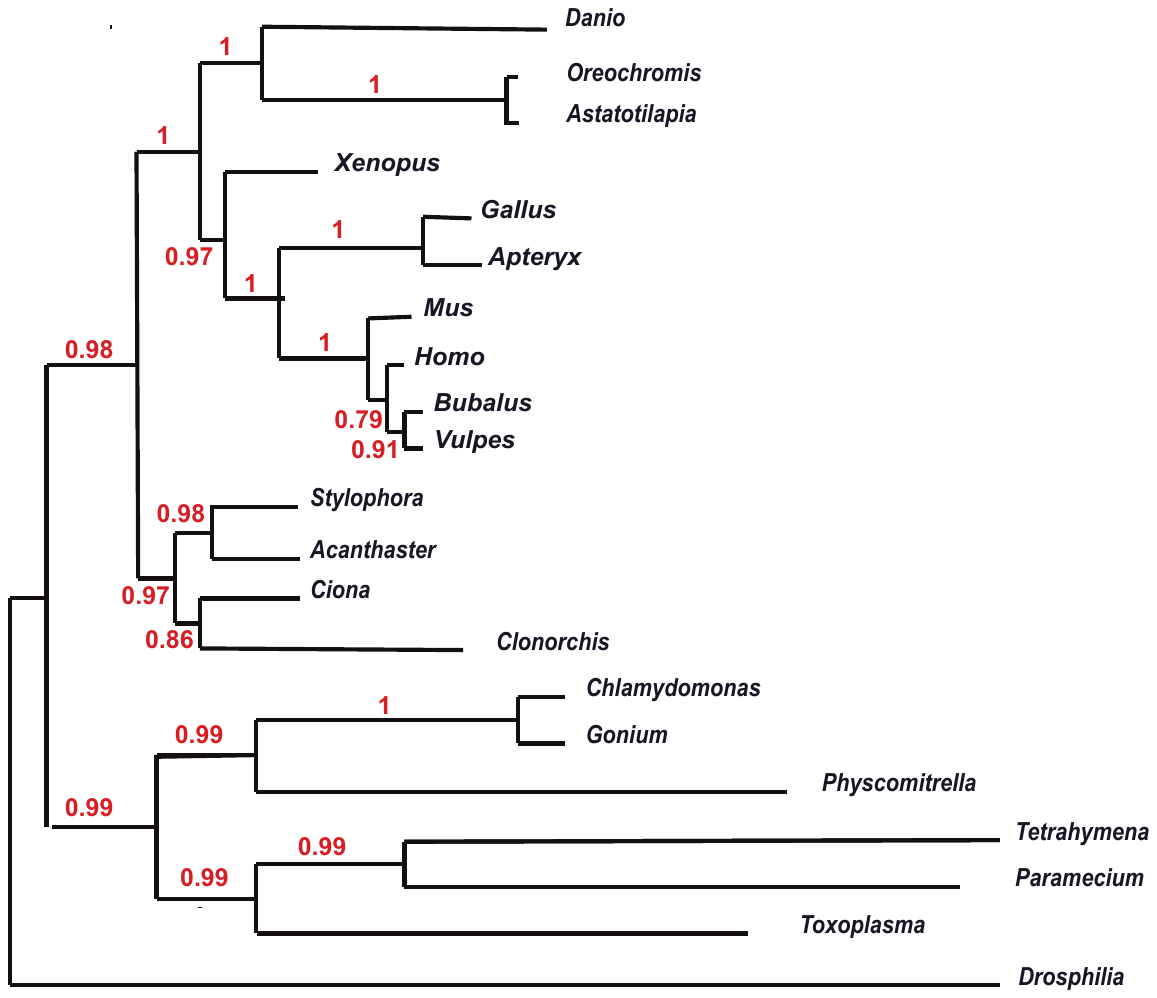
**Fig. S3. Bootstrap analysis of CFAP57**

The organisms used and their NCBI name are given below (1). Bootstrap analysis involves 1000 replicates and are shown in red (1).

**Fish:** *Danio rerio* (XP_021324203), *Oreochromis niloticus* (XP_019210353), *Astatotilapia calliptera* (XP_026024825)

**Amphians:** *Xenopus tropicallis* (XP_002931652)

**Birds:** *Gallus gallus* (XP_422397), *Apteryx rowii* **(**XP_025942666),

**Mammals:** *Mus musculus* (NP_081065), *Homo sapiens* (XP_016855911), *Bubalus bualis* (XP_006068359), *Vulpes vulpes* (XP_025847857)

**Urochordata:** *Stylophora pistillata* (XP_022792430), *Acanthaster planci* (XP_022105499), *Ciona intestinali*s (XP_018671994)

**Trematode:** *Clonorchis sinensi* (RJW62559)

**Green algae:** *Chlamydomonas reinhardtii* (PNW83807), *Goium pectoral* KXZ43455, **Moss:** *Physcomitrella patens* (XP_024357376)

**Ciliates:** *Tetrahymena thermophila* (XP_00125717), *Parmecium aurelia* (XP_001459024),

**Apicomplexa:** *Toxoplamsa gondii* (KFG58296),

**Insects:** *Drosophila melanogaster* (NP_611687)

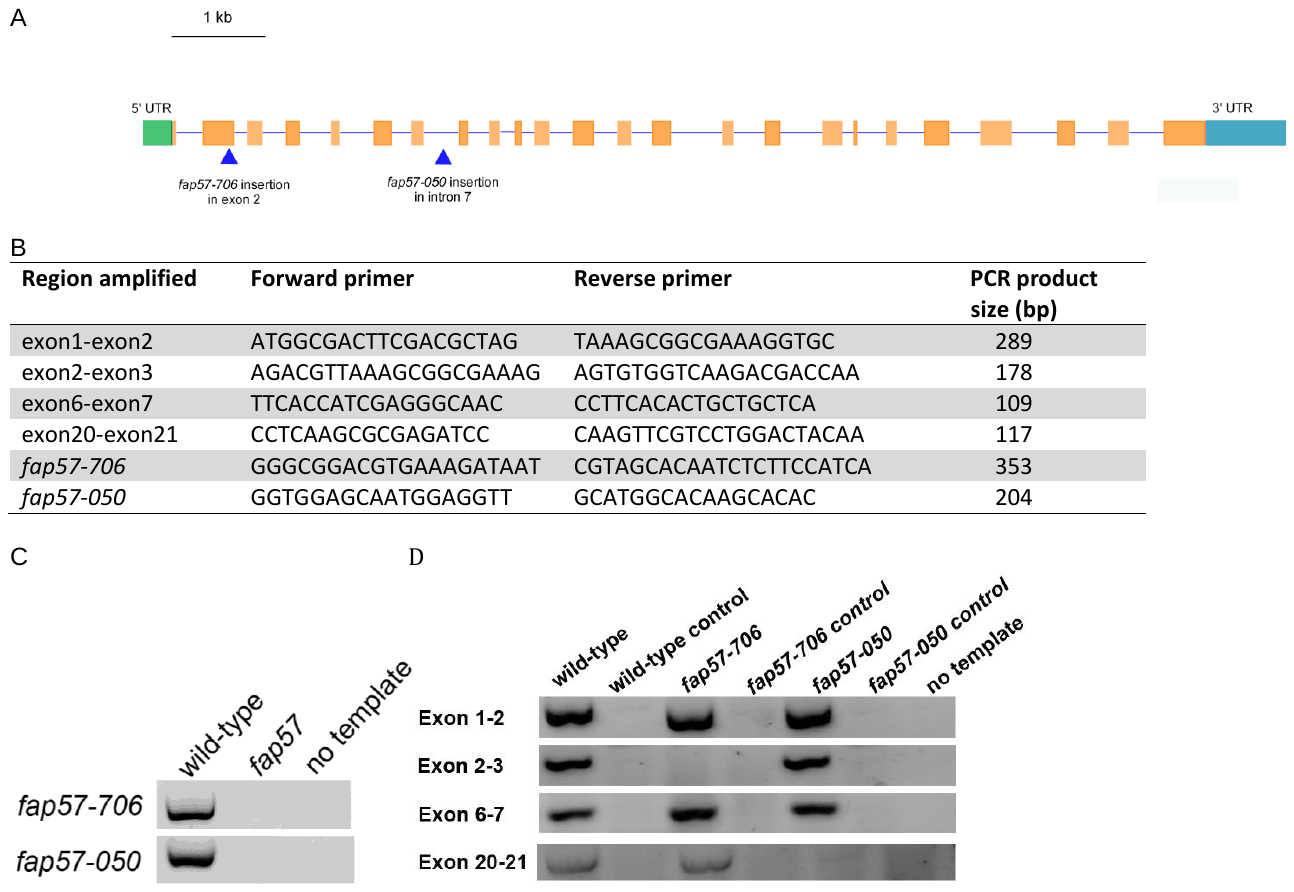
**Fig. S4. Molecular analysis of insertional mutants in *Chlamydomonas.***

(A) Diagram of *FAP57* genomic DNA with arrows showing the location of the *fap57-050* and *fap57-706* insertions. (B) List of primers used to amplify *CFAP5*7 cDNA and genomic DNA.(C) Locations verified for the LMJ.RY0402.157050 and LMJ.RY0402.107706 insertions by PCR. (D) PCR amplification of regions of *fap57* cDNA. No template indicates that no DNA was added into the reaction.

**Fig. S5. Ciliary waveform analysis in *fap57* *Chlamydomonas*.**

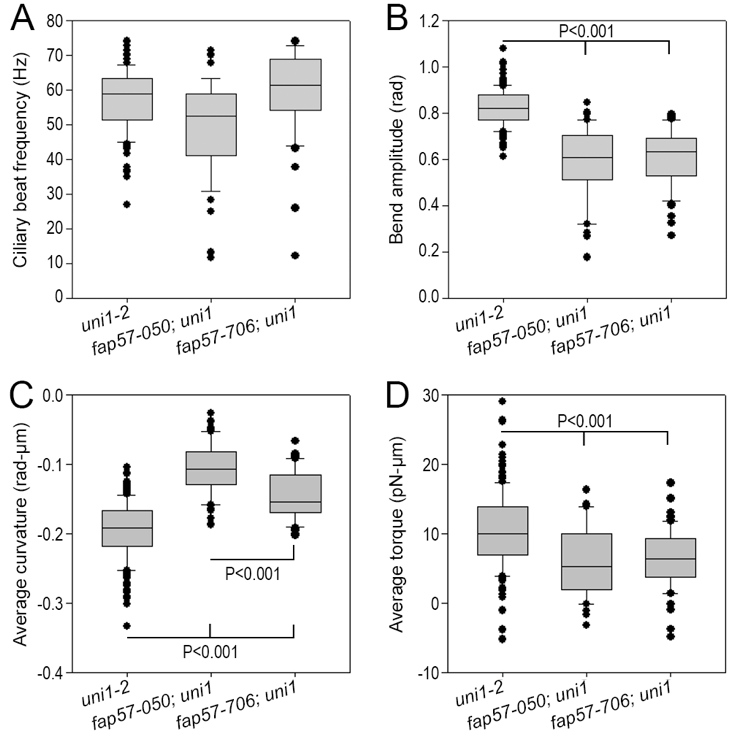

(A) Ciliary beat frequency (Hz) estimated from tangent angle. (B) Bend amplitude (rad). (C) Average curvature (rad/µm). (D) Average torque applied by the cilium about the center of the cell body (pN-µm).

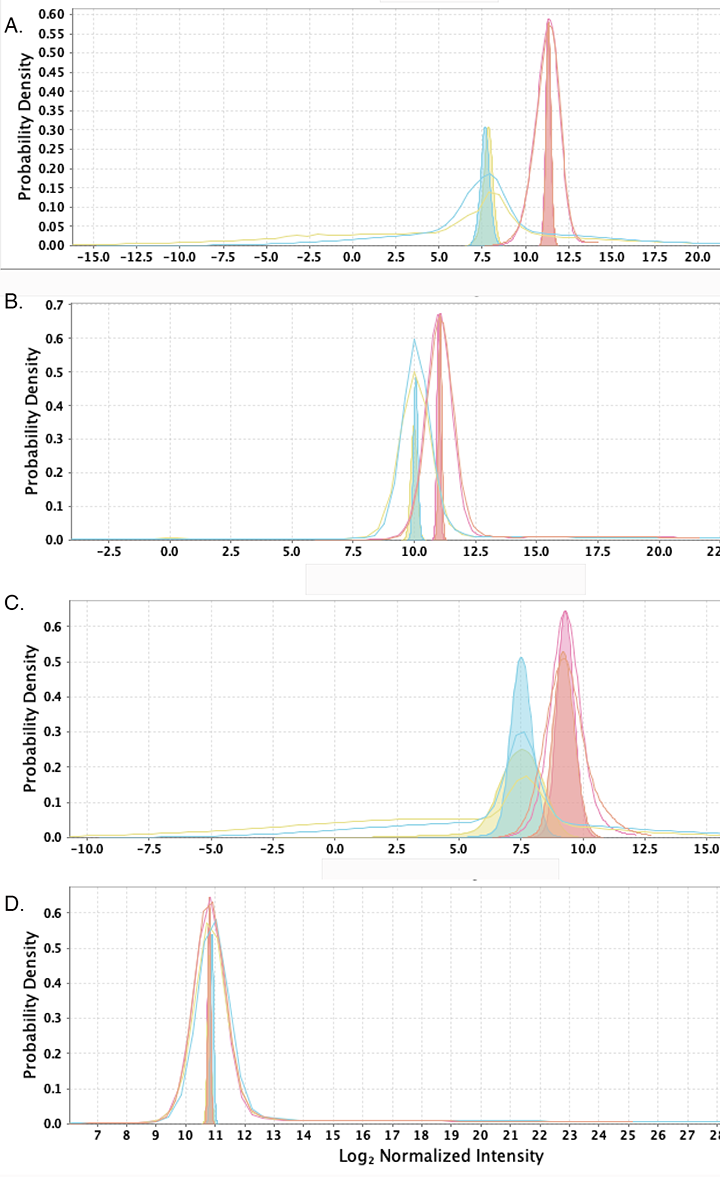
**Fig. S6. Density plots of four proteins from TMT mass spectroscopy of wild-type and mutant *Chlamydomonas*.**

**(**A). Comparison of peptides from wild-type and mutant peptides for FAP57 (Cre04.g217917) . (B). Comparison of peptides from wild-type and mutant peptides for DHC7 (Cre06.g265950). (C). Comparison of peptides from wild-type and mutant peptides for WDR49 paralog (Cre13.g562800). (D). Comparison of peptides from wild-type and mutant peptides for DHC1; I1/f dynein (Cre12.g48425). Orange: CC-125; Pink CC-4533 (parent of the insertional mutants); Yellow: *fap57-050*; Blue: *fap57-705.* Panels A, B, and C show significant differences between the wild-type and mutant samples.

Table S1. List of the top 10 genes identified in the proband using PSAP analysis.

| **Gene** | **Transcript** | **Position (hg19)** | | **Type** | **AA Change** | **exAC Freq** | **CADD** | **PSAP pvalue** |
| --- | --- | --- | --- | --- | --- | --- | --- | --- |
| *CD2* | ENST00000369478.3 | 1:117311391_C>T | nonsynonymous SNV | | P348S | 0.0015 | 17.22 | 2.00E-06 |
| *WDR65/CFAP57* | ENST00000372492.4 | 1:43675420_C>T | stopgain | | R588X | 1.72E-05 | 37 | 3.00E-06 |
| *ETS1* | ENST00000392668.4 | 11:128350323_A>T | nonsynonymous SNV | | F296I | 3.00E-04 | 25.7 | 5.00E-06 |
| *TMOD4* | ENST00000416280.2 | 1:151143016_T>C | nonsynonymous SNV | | M263V | 0.0046 | 13.71 | 1.20E-05 |
| *TEX15* | ENST00000256246.2 | 8:30695552_C>T | nonsynonymous SNV | | E2367K | 9.19E-05 | 16.87 | 1.30E-05 |
| *CEND1* | ENST00000330106.4 | 11:788386_G>C | nonsynonymous SNV | | S64W | 2.60E-05 | 10.56 | 1.80E-05 |
| *CTU2* | ENST00000453996.2 | 16:88780568_C>T | nonsynonymous SNV | | R344W | 1.67E-05 | 18.85 | 2.00E-05 |
| *HSD3B1* | ENST00000528909.1 | 1:120054192_G>T | nonsynonymous SNV | | R71I | 0.0039 | 11.87 | 2.50E-05 |
| *CCDC74B* | ENST00000310463.6 | 2:130902418_A>G | nonsynonymous SNV | | L51P | 1.00E-04 | 13.63 | 2.70E-05 |
| *GUCY2F* | ENST00000218006.2 | X:108638614_C>T | nonsynonymous SNV | | E794K | 0.0042 | 27.4 | 5.70E-05 |

**Supplemental Table 1**. List of the top 10 genes identified in the proband using PSAP analysis. The “PSAP” p-value is a measure of significance or “surprise” of sampling a particular genotype. It represents the probability of sampling a genotype given a population genetic model trained on a large database of “normal” control individuals. Smaller p-values are more surprising or unlikely (2, 3)

**Table S2.** **Measurements of wild-type and mutant *Chlamydomonas***

|  | ***uni1-2*** | ***fap57-050; uni1-2*** | ***fap57-706; uni1-2*** |
| --- | --- | --- | --- |
| **Body rotation**  **Rotations/sec** | 2.6 ± 1.0 | 1.0 ± 0.6 | 1.0 ± 0.6 |
| **Beat frequency (Hz)** | 57.4 ± 8.6 | 49.6 ± 13.5 | 59.1 ± 12.5 |
| **Bend amplitude**  **(rad/µm)** | 0.82 ± 0.08 | 0.59 ± 0.16 | 0.61 ± 0.12 |
| **Average curvature**  **rad/µm** | -0.20 ± 0.04 | -0.11 ± 0.04 | -0.14 ± 0.04 |
| **Force**  **(pN/µm)** | 10.5 ± 5.8 | 5.8 ± 5.6 | 6.3 ± 4.3 |
| **Internal forces**  **(pN/µm)** | 193 ± 42 | 188 ± 56 | 187 ± 53 |

**Table S3. TMT values from mass spectroscopy of isolated axonemes**

|  |  | ***Log_2_ Fold Change*** | | |  |
| --- | --- | --- | --- | --- | --- |
| **Protein/Gene** | **No.**  **Peptides** | **Wild-type**  **Average of wild-type**  **(n=4)** | ***fap57-1***  **replicates** | ***fap57-2***  **replicates** | ***Quantitized***  ***Spectral Count*** |
| **Dynein arms that are reduced in mutants** | | | | | |
| DHC7; **g**  Cre06.g265950 | 341 | -0.025 | -0.9/-1.1 | -0.8/-1.3 | 222 |
| DHC2; **d**  Cre09.g392282 | 540 | 0 | -0.3/-0.4 | -0.4/-0.4 | 358 |
| DHC3 (minor IDA)  Cre06.g265950 | 78 | -0.075 | -1.0/-1.0 | -0.9/-1.1 | 43 |
| **Dynein arms that are not reduced in *fap57* mutants** | | | | | |
| ODA11; DHC13; α  Cre11.g476050 | 1662 | 0.025 | 0.1/0.2 | 0.1/0 | 1124 |
| ODA4; DHC14; β  Cre09.g403800 | 1471 | 0 | 0.2/0.2 | 0.1/0.1 | 490 |
| ODA2; DHC15; γ  Cre01.g058400 | 1443 | 0.025 | 0.1/0.2 | 0.1/0 | 345 |
| DHC6; **a**  Cre05.g244250 | 444 | 0.05 | 0.2/0.2 | 0.2/-0.1 | 285 |
| DHC5; **b**  Cre02.g107050 | 342 | 0 | 0.2/0.2 | 0.1/0 | 202 |
| DHC9; **c**  Cre02.g141606 | 609 | -0.05 | 0.1/0.1 | 0/0 | 199 |
| DHC8; **e**  Cre16.g685450 | 346 | -0.025 | 0.2/0.2 | 0.1/0 | 211 |
| DHC1; **I1/f α**  Cre12.g484250 | 686 | 0 | 0.1/0.2 | 0/0 | 475 |

**Table S4: Suppression of the motility defect of *pf10***

| **Genotype** | **Ratio of cells in supernatant to total cell number**  **(n= 300)** |
| --- | --- |
| *FAP57; pf10* | 0.1 |
| *FAP57; PF10* | 0.97 |
| *fap57-050; pf10* | 0.94 |
| *fap57-706; pf10* | 0.95 |

**Table S5. Antibodies used in this study**

| **Antibody** | **Antigen** | **Source** | **Dilution**  **Immunoblot** | **Dilution**  **IF** |
| --- | --- | --- | --- | --- |
| HPA028623 rabbit polyclonal | CFAP57 | Sigma-Aldrich,  St. Louis, MO | 1:1000 | 1:500 |
| HPA036225 rabbit polyclonal | WDR49 | Sigma-Aldrich,  St. Louis, MO | 1:500 | 1:200 |
| 6-11B-1, monoclonal  T7451 | Acetylated -tubulin | Sigma-Aldrich,  St. Louis, MO | NT | 1:1000 |
| HPA053129  rabbit polyclonal | DNALI1 | Sigma-Aldrich,  St. Louis, MO | NT | 1:500 |
| 5600 | RSPH1 | Abmart, Berkeley Heights, NJ | NT | 1:250 |
| Mouse monoclonal | DNAI1 | University of North Carolina | NT |  |
| Alexa Fluor-488 Secondary antibody |  | Life Technologies, Carlsbad, CA |  |  |
| Alexa Fluor-647  Secondary antibody |  | Life Technologies,  Carlsbad, CA |  |  |
| indocarbocyanine (CY3 conjugated secondary antibody |  | Jackson ImmunoResearch Laboratories, West Grove, PA |  |  |
| Rhodamine Red-X (RRX) conjugated secondary antibody |  | Jackson ImmunoResearch Laboratories, West Grove, PA |  |  |

**Table S6. Primer sequences for analysis of *CFAP57***

| **Primer used for amplification of**  **genomic DNA** | | |
| --- | --- | --- |
| **Position** | **Primer Sequence (5’ to 3’)** | **Name** |
| Exon 11 | GAT GCC AAA AGG GGA TAG A | M13 tag sense primer |
|  | CAG GGC TAT GGC TCC TTT C | M13 tag antisense primer |
| **Primers used for amplification of cDNA** | | |
| **Gene** | **Primer Sequences (5’ to 3’)** | **PCR product size (bp)** |
| *CFAP57* | GGG GAG AAC CCC CAA CCA TAT CT  TGA TGG GTG CTG AGT GCA AT | 597 |
| *DNAI1* | AGA GAA GGA GAA GGC AAA GAC CCC  TGT ACT CAG GGA AGC TGG GGT TCT | 400 |
| *PPIA* | CCG TGT TCT TCGACA TTG CC  ACA CCACAT GCT TGC CAT CC | 371 |

Movie S1-S6 (separate file).

S1: Video recording of control human nasal cells in profile demonstrating normal waveform.

Video was recorded at 200 fps using a 60x objective with DIC optics. Playback speed is 15% of

normal speed. Scale bar, 4 μm.

S2: Video recording of PCD 2-II human nasal cells in profile demonstrating heterogeneous

waveform. Video was recorded at 200 fps using a 60x objective with DIC optics. Playback speed

is 25% of normal speed. Scale bar, 5 μm.

S3-S4: Control hTEC transduced with a non-targeted shRNA sequence. The movie

shows the normal ciliary waveform, at normal speed (S1) and slowed to 1/10x (S2).

S5-S6: Representative images of hTEC transduced with a targeted CFAP57 sequence

(sequence 3). The movie shows shows a reduced curvature, at normal speed(S3) and slowed to

1/10x (S4).

**References**

1.	Dereeper A*, et al.* (2008) Phylogeny.fr: robust phylogenetic analysis for the non-specialist. *Nucleic acids research* 36(Web Server issue):W465-469.

2.	Karczewski KJ*, et al.* (2017) The ExAC browser: displaying reference data information from over 60 000 exomes. *Nucleic acids research* 45(D1):D840-D845.

3.	Rentzsch P, Witten D, Cooper GM, Shendure J, & Kircher M (2019) CADD: predicting the deleteriousness of variants throughout the human genome. *Nucleic acids research* 47(D1):D886-D894.
