## Supplementary material for "Mutation of *CFAP57* causes primary ciliary dyskinesia by disrupting the asymmetric targeting of a subset of ciliary inner dynein arms": Legends for movies

Supplemental movie

S1-S2: Control hTEC transduced with a non-targeted shRNA sequence. The movie shows the normal ciliary waveform, at normal speed (S1) and slowed to 1/10x (S2).

S3-S4: Representative images of hTEC transduced with a targeted *CFAP57* sequence (sequence 3). The movie shows shows a reduced curvature, at normal speed(S3) and slowed to 1/10x (S4).
